# Outbreak dynamics of high pathogenicity avian influenza virus H5N1, clade 2.3.4.4b euBB, in black-headed gulls and common terns in Germany in 2023

**DOI:** 10.1101/2025.09.17.676714

**Authors:** Ulrich Knief, Ann Kathrin Ahrens, Valerie Allendorf, Carla J. Behringer, Justine Bertram, Sandra Bouwhuis, Wolfgang Fiedler, Anja Globig, Anne Günther, Christof Herrmann, Sascha Knauf, Dominik Marchowski, Simon Piro, Anne Pohlmann, Robert E. Rollins, Christoph Staubach, Timm Harder

## Abstract

Since winter 2022/23, high pathogenicity avian influenza virus (HPAIV) H5N1 clade 2.3.4.4b, genotype euBB, has caused extensive mortality among wild birds. This genotype emerged in France in spring 2022 through reassortment between a gull-adapted low-pathogenicity virus and HPAIV H5N1. Phylogeographic and spatiotemporal analyses show that transmission into German breeding colonies of black-headed gulls (*Chroicocephalus ridibundus*) and common terns (*Sterna hirundo*) involved multiple independent incursions, likely via black-headed gulls returning from wintering grounds in the Netherlands, Belgium, and France. It spilled over into common terns, which breed in shared colonies with black-headed gulls, and led to high adult mortality in both species in 2023 (at least 8,137 black-headed gulls and 614 common terns; >3% of the breeding population), followed by significant breeding pair declines in 2024 (−15.9% in black-headed gulls, −5.8% in common terns). Increased immunity, at least in common terns, may have contributed to the apparent fade-out of genotype euBB. These findings highlight how integrating ornithological, epidemiological, and virological data can aid our understanding of viral transmission routes and population-level impacts, while also stressing that HPAIV should be added to the growing list of pressures on seabirds, a group that was already the most threatened among all bird taxa globally.

**Research Highlights:**

- HPAIV H5N1 euBB caused mass mortality in gulls and terns in Germany in 2023
- Multiple independent viral incursions occurred into different parts of Germany
- Serology showed survival and rising H5-seroprevalence in common terns
- Outbreaks reduced the black-headed gull breeding populations by up to 24%

## 1. Introduction

Since their initial detection in 1996 in a flock of geese in southern China, high pathogenicity avian influenza H5Nx viruses (HPAIV) of the so-called goose/Guangdong lineage (gs/GD) have continued to evolve to form up to ten main phylogenetic lineages (clades), with further phylogenetic divergence into genetic subclades (Xie et al., 2023). Frequent bidirectional spillover between wild bird populations and poultry has created opportunities for co-infection and reassortment with low pathogenicity (LP) AIV, leading to the emergence of numerous genotypes (Fusaro et al., 2024; King et al., 2022). Viruses of subtype H5N1 of the fifth order clade 2.3.4.4b have spread globally in successive pan-zoonotic dissemination waves since 2020.

The widespread presence of infected sick and dead wild birds in the environment has led to an increased incidence of scavenger-borne infections in wild carnivorous mammals and in the fur industry (summarized in ENETWILD Consortium et al., 2024; Kareinen et al., 2024), and in domestic cats (Rabalski et al., 2023). In the vast majority of cases, mammals, including humans, have proven to be dead end hosts for these viruses (Peacock et al., 2025). In contrast, HPAIV H5Nx-infected wild birds, particularly those of colony-breeding seabird species, have experienced mass mortalities on numerous occasions (Knief et al., 2025; Knief et al., 2024; Pohlmann et al., 2023), threatening the persistence of these often already highly endangered species.

The variability of viruses from the gs/GD lineage, in particular of clade 2.3.4.4b, is largely due to their ability to reassort with co-circulating LPAIV (King et al., 2021). Co-infection of an avian host provides the opportunity for exchange of the octo-segmented genome between two parental influenza viruses. Each of these novel combinations (i.e., reassortants) may exhibit novel phenotypic properties that affect the fitness of these viruses and give rise to epidemiological advantages or disadvantages (Piesche et al., 2024). The propensity for reassortment of the currently circulating clade 2.3.4.4b seems to be particularly pronounced, which may also be associated with its extensive, nearly global spread (Kandeil et al., 2023).

AIV generally display a lack of selectivity with regard to their avian hosts (Klaassen & Wille, 2023). Nevertheless, a few subtypes are known to circulate predominantly within specific avian groups, i.e. Anseriformes or Charadriiformes. The subtypes H13 and H16, for example, are predominantly found in Charadriiformes, particularly in Laridae, but are detected only rarely in Anseriformes, indicating an adaptation at the host species level (Verhagen et al., 2014). In contrast, viruses of the gs/GD lineage H5Nx have been detected in almost all exposed avian taxa, with the current exception of some species in the Columbidae family, which are considered to be largely refractory to infection (Klaassen & Wille, 2023).

In the early summer of 2022, a novel reassortant between the gs/GD clade 2.3.4.4b and an H13 virus emerged (Briand et al., 2025; Fusaro et al., 2024). This reassortant was identified on the west coast of France and is now designated as genotype euBB. The spread of this genotype was rapid and massive across Europe and affected poultry, as well as a large number of wild avian species. Among those, both black-headed gulls (*Chroicocephalus ridibundus*) and common terns (*Sterna hirundo*) were the major hosts of genotype euBB and dominated its epidemiology in Europe as a whole, and in Germany in particular (Fusaro et al., 2024; Indykiewicz et al., 2025). In this paper, we focus on these two species and describe how the HPAIV H5N1 genotype euBB spread and affected populations in Germany.

## 2. Material and Methods

### 2.1. Spatiotemporal analysis of HPAIV detection and prevalence in black-headed gulls and common terns

To evaluate changes in HPAIV prevalence over time, we retrospectively analysed data from the Database for Avian Influenza in Wild Birds (Aviäre Influenza Wildvogel-Datenbank, AI-DB), Animal Disease Information System (ADIS) of the European Union, WAHIS database and the German National Database for Animal Diseases (Tierseuchennachrichtensystem, TSN) dating back to 2006. While the TSN usually reports sick, killed or dead birds infected with HPAIV H5, AI-DB and ADIS also contain data from active surveillance.

### 2.2. Quantitative HPAIV-related mortality data in breeding colonies in 2023

For the 2023 breeding season, we collected data on German colonies of black-headed gulls and common terns by reaching out to local administrators, site managers, NGOs, scientists, and other stakeholders. We also issued a call for collaboration through newsletters distributed by the German ringing schemes and the German Ornithological Society (Deutsche Ornithologische Gesellschaft, DOG). This form of data collection relied on voluntary responses. Given that there is no legal obligation to report (mass) mortality events, nor a current national overview of all breeding colonies (see below), we cannot guarantee completeness of the dataset. In total, however, we gathered HPAIV data from 52 black-headed gull and 48 common tern colonies (**Supplementary Data**).

For each colony, we determined whether it was affected by HPAIV. A colony was classified as affected if adults or juveniles either tested positive for HPAIV, exhibited unusual mortality, or displayed neurological clinical signs suggestive of HPAIV infection. Specifically, we requested data on the location of the colony, the affected species and age classes (juveniles or adults), the number of breeding pairs, and the total number of individuals found dead, categorized by age class and species. For about half of the colonies (*N* = 31 black-headed gull and *N* = 21 common tern colonies), we received the date of the first fatality (defined as the onset of infection in a colony). We also recorded whether the dead birds tested positive for HPAI, and if so, the specific virus type (see above). For a subset of colonies (*N* = 9 black-headed gull and *N* = 6 common tern colonies), we received longitudinal, i.e. within-season repeated, data, which allowed us to reconstruct fatality curves representing the cumulative number of adult or juvenile birds found dead or dying over time. To estimate breeding bird densities in these 15 colonies, we also gathered data on the estimated size of the colony area.

### 2.3. Data for quantifying the impact of HPAIV-related mortality

#### 2.3.1. Quantifying excess mortality from black-headed gull ring recovery data 2000– 2023

For the period 2000–2022, we extracted data on recoveries of dead black-headed gulls and common terns from the database of the Hiddensee bird ringing scheme (covering eastern Germany). Using these data, we calculated the average number of ringed birds reported dead per month to estimate what we call background mortality. We then compared these baseline numbers with the monthly dead recoveries recorded in 2023. Recoveries categorized as “only ring found” or “only colour mark found”, as well as those of chicks that had died before fledging, were excluded from the analysis.

#### 2.3.2. Breeding bird census data 2022–2024

In context of the reporting requirements under article 12 of the EU Birds Directive for the period from 2012 to 2016, the black-headed gull and common tern breeding populations were estimated at 115,000–160,000 (2016) and 8,500–9,000 (2012–2016) breeding pairs, respectively (Ryslavy et al., 2020). The most recent nationwide data on the geographic breeding distribution of both species in Germany were collected between 2005 and 2009 (Gedeon et al., 2014). We use these data to visualize the breeding ranges of the two species. We obtained current breeding pair counts for both species from the Federation of German Avifaunists (DDA) that were collected as part of their *Gulls & Terns* monitoring protocol in frame of the *German Rare Breeding Bird* scheme (Wahl et al., 2020). This dataset spans the years 2022–2024 and we included only colonies consistently surveyed across all three years (*N* = 177 colonies). These data represent around 10% of the estimated national breeding population from 2016 for both species (Ryslavy et al., 2020).

For a more comprehensive analysis, we compiled an additional dataset for 2022–2024 that incorporates data from several entire German federal states, national parks and from the Szczecin Lagoon in Poland (**Supplementary Data**). The inclusion of the Polish Szczecin Lagoon was necessary because of a strong movement of breeding pairs of both species from German colonies in Mecklenburg-Western Pomerania to two recently created artificial bird islands in Polish waters (Marchowski et al., 2024). This dataset was combined with the DDA data. For 2022, the combined dataset included 89,482 breeding pairs of black-headed gulls (with 705 pairs from the Szczecin Lagoon) and 7,645 breeding pairs of common terns (with 157 pairs from the Szczecin Lagoon), representing approximately 55.5–77.2% and 83.2–88.1% of the estimated 2012–2016 German populations, respectively. For common terns, the dataset likely captured nearly all known breeding pairs in 2022 and subsequent years, suggesting that the observed decrease of over 10% compared to 2016 represents a true population decline rather than an artifact of incomplete data. For black-headed gulls, the absence of data from several federal states renders any conclusion about a true population decline between 2016 and 2022 more speculative.

Black-headed gull and almost all common tern colonies were affected by HPAIV for the first time in the breeding season 2023. Therefore, we use 2022 as the reference year for the population size prior to any HPAIV outbreaks and interpret the declines in 2023 and 2024 as a direct consequence of HPAIV, as such drastic population reductions are unlikely to be caused by other factors.

### 2.4. Data on black-headed gull wintering areas and their overlap with HPAIV outbreaks

Ringing data were available from the federal states of Mecklenburg-Western Pomerania and Saxony (bird ringing scheme Vogelwarte Hiddensee), Schleswig-Holstein, Lower Saxony, and North Rhine-Westphalia (bird ringing scheme Vogelwarte Helgoland). For other federal states, ringing data were either unavailable or available in very low numbers due to a lack of or very low gull ringing activity.

We selected ring recovery data based on the following criteria:

(i) Known origin: We included only birds ringed as nestlings or recorded as adults in breeding colonies during the breeding season (1 April to 31 July). Adult birds were either caught on the nest or identified from a distance via colour rings.
(ii) Unique recoveries: For birds with multiple records at the same wintering location in the same winter season, we retained only one recovery per winter season but included recoveries from different seasons.
(iii) Time period: Records from 2000 to 2024 were included.
(iv) Winter definition: The winter period was defined as 16 November to 15 February.

After applying these criteria, we retained 1894 recoveries distributed across the five federal states: Mecklenburg-Western Pomerania (*N* = 1119), Schleswig-Holstein (*N* = 255), Lower Saxony (*N* = 231), Saxony (*N* = 200), and North Rhine-Westphalia (*N* = 89).

### 2.5. Sequencing and phylogeographic analyses

Sequencing of avian influenza-positive samples was performed using an amplicon-based protocol on nanopore platforms (Oxford Nanopore Technology, Oxford, UK). Briefly, RNA extracted using commercial silica matrix kits was transcribed into DNA using the Superscript III One-Step and Platinum Taq kit (#12574026, Thermo Fisher Scientific, USA) with influenza-specific primers (Pan-IVA-1F_BsmF: TATTCGTCTCAGGGAGCRAAAGCAGG; Pan-IVA-1R_BsmR: ATATCGTCTCGTATTAGTAGAAACAAGG), each binding to the conserved 3’ or 5’ end of all influenza RNA segments. DNA amplicons were purified with Agencourt AMPure XP magnetic beads (#A63881, Beckmann Coulter, Krefeld, Germany) using DNA LoBind Tubes (#0030108051, Eppendorf, Wesseling-Berzdorf, Germany). Quantification of nucleic acids was done with Qubit Fluorometry (Thermo Scientific). Approximately 400 ng of cDNA was sequenced using a transposase-based library preparation approach with Rapid Barcoding (SQK-RBK114, Oxford Nanopore Technologies, Oxford, UK) and on a MinION Mk1B or PrometION P2 Solo instrument with the latest MinKNOW software core (v6.0.15). High accuracy base calling of the raw data using Dorado (v7.4.14, Oxford Nanopore Technologies) was followed by demultiplexing, a quality check and a trimming step to remove poor quality, adapter and short (<20 bp) sequences. The generated data were stored in FASTQ and POD5 data formats. The bioinformatics software suite Geneious Prime (GraphPad Software LLC, version 2024.0.1) was used for analysis. Sequences were trimmed to remove primer sequences. Consensus sequences were obtained using an iterative map-to-reference approach with Minimap2 (vs 2.17; Li, 2021). Reference genomes were selected from a curated collection of all HA and NA subtypes and a selection of internal gene sequences to cover all potentially circulating viral strains. Polishing of the final genome sequences and annotation were performed manually after consensus generation (threshold matching 60% of bases of total adjusted quality). Segment-specific and concatenated whole-genome multiple alignments were generated with MAFFT (v7.450) and subsequent maximum likelihood (ML) trees were calculated with RAxML (v8.2.11; Stamatakis, 2014) using a GTR GAMMA model selected by use of ModelTest with rapid bootstrapping and search for the best scoring ML tree, supported by 1000 bootstrap replicates, or alternatively with FastTree (v2.1.11; Price et al., 2010). Subsets of closely related full genomes (the largest open reading frames of each segment had been concatenated) were extracted and used for further ML phylogenetic analyses. Time-scaled trees of concatenated sequences of the different genotypes were calculated using the BEAST software package (v10.5.0; Suchard et al., 2018), using a GTR GAMMA substitution model, an uncorrelated relaxed clock with a lognormal distribution, and coalescent constant population tree models. Chain lengths were set to 50 million iterations and convergence was checked using Tracer (v1.7.1; Suchard et al., 2018). Time-scaled summary maximum clade credibility (MCC) trees with 10% posterior burn-in were generated using TreeAnnotator (v1.10.4; Suchard et al., 2018) and visualized using FigTree (v1.4.4; Rambaut, 2018). The MCC trees were checked for robustness by 95% highest posterior density (HPD) confidence intervals at each node and posterior confidence values as branch support. Spatiotemporal spread was inferred on MCC trees with country, states, region or continuous GPS features using SPREAD (v1.0.7; Bielejec et al., 2011) and visualized using QGIS (v3.34.13; QGis.org, 2024). Centroids were used for latitude and longitude information for countries and states. For detailed geocoordinates of each sequence, latitude and longitude were extracted using metadata and adapted optical means.

### 2.6. Active surveillance for HPAIV in black-headed gulls and common terns in northern Germany

#### 2.6.1. Black-headed gulls in Mecklenburg-Western Pomerania, Germany

The nature reserve “Isle of Böhmke”, an island of about 0.03 km^2^ is located in the southeastern part of the Achterwasser of Usedom, northeastern Germany (53°56′42″ N, 14°01’59″ E). It is known to harbour a black-headed gull breeding colony during the months April-July. On the 30^th^ of May 2023, 25 adult individuals were caught and sampled alive, during regular bird ringing activities coordinated by local ornithologists. The sampling was performed by wildlife experienced veterinarians. The study approval was granted by the regional authority for animal experiments, the Landesamt für Landwirtschaft, Lebensmittelsicherheit und Fischerei (LALLF) under the permit number LALLF 7221.3-2-007/23.

Each bird was sampled for a combined oropharyngeal and cloacal swab (*N* = 25) which was stored in virus cultivation medium (SIGMA VIROCULT ® - virus transport medium, Medical Wire & Equipment Ltd. England & Wales). Subsequently, a corresponding blood sample (*N* = 22) was taken from the *Vena ulnaris* using a 27G-needle in combination with an attached 1ml-syringe. The collected blood was immediately transferred into a reaction tube and cooled until arrival to the laboratory. Not more than 1 ml blood was taken from each bird. The handling and sampling of animals followed good veterinary practice procedures. A single adult BHG, that was found dead on the same day within the colony, was also swab sampled. All samples were transported at +4°C to the laboratories of the Friedrich-Loeffler-Institut where they were analysed for the presence of IAV virus RNA and IAV or H5-specific antibodies, respectively.

Processing of those sample matrices was conducted as described before in Günther et al. (2024). Briefly, the swab sample supernatant represented input for automated RNA extraction, that included extraction controls and a heterologous control RNA. An initial RT-qPCR screening targeted the matrix protein gene of IAV as detailed in Hassan et al. (2022). Inactivated serum samples were screened in NP-specific, and in case of positive findings subsequently in H5-specific, commercial ELISAs as detailed in section 5.6.2.

#### 2.6.2. Common terns in Lower Saxony, Germany

A mono-specific common tern colony situated at the Banter See in Wilhelmshaven at the German North Sea coast (53°30′40″ N, 08°06’20″ E) has been systematically monitored since 1992. Since then, birds of known age and sex have been transponder-marked. As part of a standardized protocol, nests are checked three times a week to monitor reproductive performance. During the three-week incubation period of the eggs, during which incubation is shared by both members of a breeding pair, an antenna is placed around the nest for at least 24 hours to assess the identity of locally hatched breeders, and for a subset of these breeders — 152, 253 and 168 in 2022, 2023 and 2024, respectively — blood samples were collected during the incubation phase, with the help of the nest antennas and by making use of larval instars of the blood-sucking bug *Dipetalogaster maximus* placed in an artificial hollow egg with small holes along the exterior (Becker et al., 2006). Hereto, a ‘bug egg’ was placed into the nest of the target bird once it was detected incubating and left there for 20–30 minutes, during which the bug was able to suck the bird’s blood. During this time, the bird was closely observed from observation huts within the colony to prevent sample contamination from partner switches. Once the bug had successfully obtained a blood meal, the blood was extracted from the bug’s abdomen using a syringe.

Nests continued to be monitored three times a week to establish hatching, and to assess the fate of the chicks. In 2023 and 2024, 12 and 143 chicks were blood sampled at the age of twenty days by pricking their brachial vein and collecting the resulting blood drops with capillary tubes. Fresh blood of both adults and chicks was separated through centrifugation and plasma was stored at −20°C until further analysis. Plasma samples were first checked for the presence of avian influenza A (AI) antibodies using a commercially available ELISA kit (ID Screen® Influenza A Antibody Competition, Innovative Diagnostics, Grabels, France) run on a 2010 EZ Read 400 microplate reader (anthos Mikrosysteme GmbH, Friesoythe, Germany). This method tests for the presence of antibodies in plasma through the formation or inhibition of coloured dyes, which can be quantified through measuring absorbance at 450nm. The percentage of antigens outcompeted (%S/N) was calculated as the OD_Probe_/OD_NC_ x 100 where OD is the optical density at 450 nm and NC refers to the negative control included in the kit. Each plate was validated through running a positive and negative control delivered with the kit according to the standard protocol. Birds were scored as positive for general AI antibodies based on the thresholds given in the protocol (i.e., %S/N < 45%). Individuals positive for general AI antibodies were further tested for H5-specific antibodies using an additional, commercially available ELISA kit (ID Screen® Influenza H5 Antibody Competition, Innovative Diagnostics, Grabels, France) which followed the same basic protocol as the general AI ELSISA. Here any bird with %S/N < 50 was considered positive for H5-specific antibodies.

### 2.7. Statistical Analyses

Unless stated otherwise, all analyses were conducted in R (v4.4.1; R Core Team, 2024). The specific packages used for each analysis are detailed below.

#### 2.7.1. Descriptive statistics, nonparametric tests and mixed-effects models

Descriptive statistics and Fisher’s exact test were conducted within R. Exact binomial confidence intervals were derived using the binom package (v1.1-1.1; Dorai-Raj, 2022). (Generalized) linear mixed-effects models ((G)LMMs) were fitted in glmmTMB (v1.1.9; Brooks et al., 2017). We modelled the number of adult birds found dead within each colony as the dependent variable. Species (factor with two levels: black-headed gull or common tern) and the date of the first recorded fatality (in Julian days) were included as fixed effects, with colony ID fitted as a random intercept. The log-transformed number of breeding adults in each colony was included as an offset term. Due to overdispersion in the data (assessed with the performance package v0.13.0; Lüdecke et al., 2021), we used a negative binomial error distribution instead of a Poisson. Significance of the random intercept was assessed using a likelihood ratio test. Back-transformed marginal parameter estimates were derived using the emmeans package (v1.10.3; Lenth, 2024).

For both black-headed gulls and common terns, we fitted a linear mixed-effects model with the log-transformed number of breeding pairs per federal state as the dependent variable, and year (factor: 2022, 2023, 2024), colony location (factor: coastal, inland) and their interaction as fixed effects, and federal state ID as a random intercept. Back-transformed marginal parameter estimates and *P*-values for pairwise comparisons were derived using the emmeans package.

#### 2.7.2. Spatial Statistics

We fitted spatially explicit models using the sdmTMB package (v0.6.0.9004; Anderson et al., 2022), which accounts for spatial covariation between colonies by incorporating a spatial random field. To construct the random field, we used geographic coordinates in the equidistant Mercator projection and the INLA package (v24.06.27; Lindgren & Rue, 2015). Geostatistical analyses were conducted with the sf package (v1.0-16; Pebesma & Bivand, 2023), and visualizations were created using colour palettes from the viridis (v0.6.5; Garnier et al., 2024) and cartography (v3.1.4; Giraud & Lambert, 2016) packages.

To model the spatial and temporal dynamics of HPAIV outbreaks across Germany, for each affected colony, we converted the date of the first fatality into Julian days and used this as our dependent variable in a spatially explicit model. We fitted species (factor with two levels: black-headed gull or common tern), age class (adult or chick) and latitude (covariate) as fixed effects. Age class was not significant and we removed it from the final model. Model fit was assessed using built-in functions from sdmTMB, visual inspection of residual distributions, and the DHARMa package (v0.4.6; Hartig, 2022). We used this model to predict the Julian day of the first fatality for all affected colonies. Based on these predicted dates, we generated equal-day contour lines to visualize the spatial and temporal spread of HPAI outbreaks across Germany. These contour lines were manually curated for illustrative purposes.

The trajectories of HPAIV outbreak waves varied considerably in both magnitude (total number of fatalities) and fatality rate increase between colonies. To investigate differences between trajectories, we used within-season longitudinal data from 12 colonies and fitted logistic curves to the cumulative number of dead adults and chicks collected in a colony over time. For each colony, species, and age class, we estimated two key parameters from the logistic curves: the asymptote and the slope. In total, we fitted *N* = 18 models across all colonies, species (*N* = 10 black-headed gull and *N* = 8 common tern models) and age classes. The asymptote represents the maximum number of fatalities reached (magnitude of the outbreak), while the slope at the inflection point indicates the rate of increase in fatalities (fatality rate increase). These two parameters were then used in subsequent linear models to assess whether the magnitude (asymptote) or the fatality rate increase (slope) of outbreaks were influenced by the breeding pair density in a colony (number of breeding pairs per colony area).

#### 2.7.3. Wintering Area Identification, Ring Recovery Mapping and Overlay of HPAI Cases

The wintering areas of German black-headed gulls were identified using the ring recovery data in QGIS (QGis.org, 2024) and the “heatmap” symbology with a radius of 100 km. A final combined map was produced to display the overlap between wintering areas and HPAI outbreak records, incorporating data from all regions.

## 3. Results

### 3.1. Passive (HP)AIV surveillance in black-headed gulls and common terns in Germany since 2022

In Germany, HPAIV spread among colonially breeding seabirds in the summer of 2022, primarily affecting terns and gannets, but also impacting gulls, leading to increased mortality (Pohlmann et al., 2023). Between 2022 and the end of 2024, data from the national Avian Influenza Database (AI-DB) in Germany showed that 804 black-headed gulls and 224 common terns were tested for HPAIV. The majority of samples was obtained through passive surveillance, with dead or diseased birds providing 85% of the samples for both species. Among black-headed gulls, 21% of the samples tested positive for HPAIV H5 in 2022, rising sharply to 53% in 2023. For common terns, 26% of samples tested positive in 2022, increasing to 36% in 2023, marking 2023 as the year with the highest positivity rate. By 2024, the situation had changed substantially. Among black-headed gulls, only one out of 85 samples tested positive for HPAIV H5. In common terns, testing was limited to 6 individuals, all of which tested negative for HPAIV H5 (**Figure 1**, **Supplementary Table S1**).

**Figure 1:**
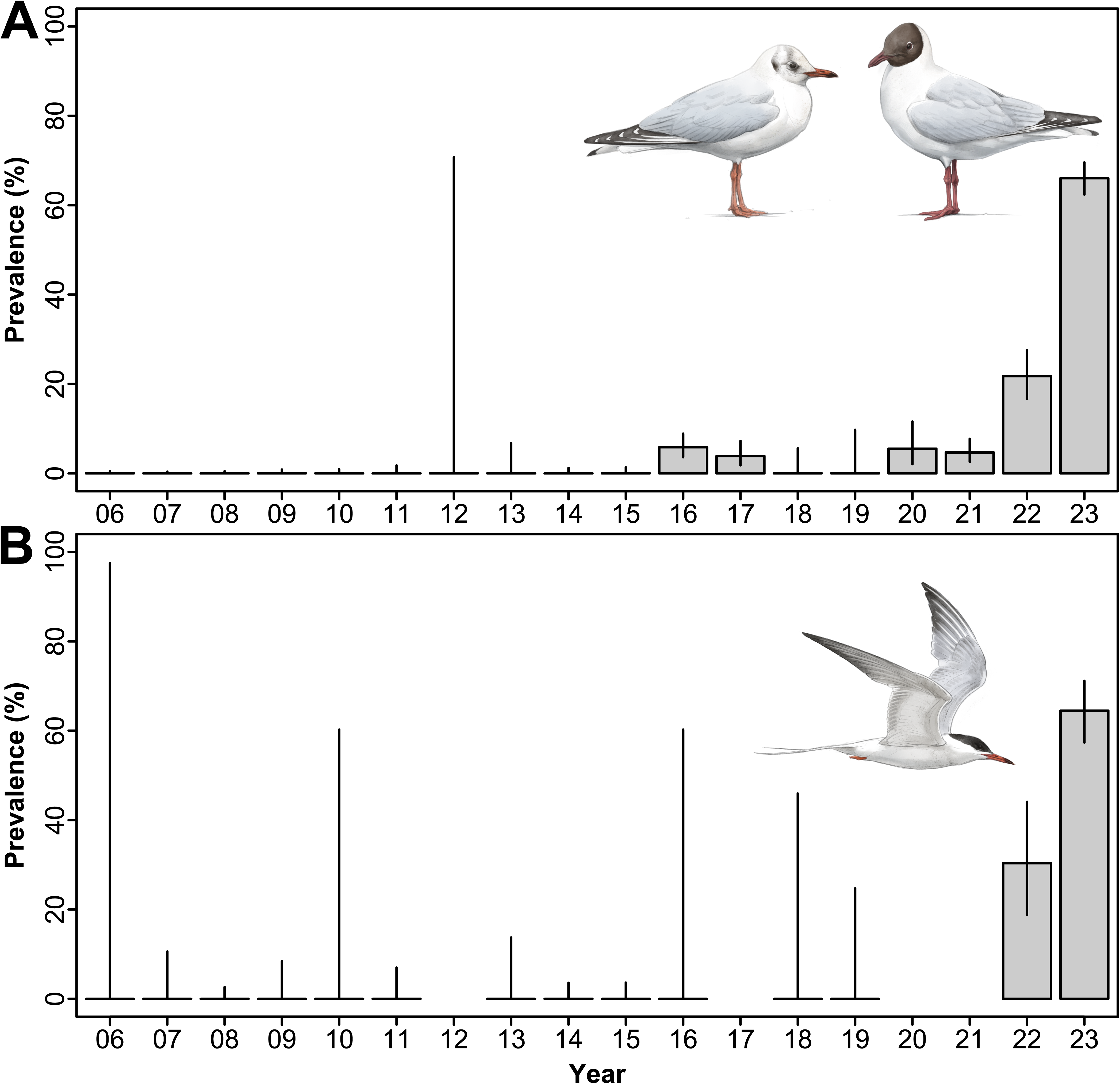
Prevalence estimates (in percent) and exact 95% binomial confidence intervals of high pathogenicity avian influenza H5 virus cases reported for (A) black-headed gull and (B) common tern samples collected in Germany between 2006 and the end of 2023. Each bar represents data gathered between 1 January and 31 December of the year shown. The bird illustrations were created by Javier Lazaro.

While these data reveal the overall trend of a major HPAIV H5N1 outbreak in 2023, they lack detailed information on the actual spread, mortality, and impact of the event. In the following sections, we first describe the 2022/23 spread of HPAIV H5N1 from the wintering grounds of black-headed gulls to Germany. Next, we quantify the mortality in German breeding colonies of black-headed gulls and common terns in 2023. Finally, we estimate the impact of the outbreak on breeding pair numbers of both species in 2024. We integrate field (ornithological) and laboratory (genetic and serological) data, which provides a more comprehensive understanding of the infection dynamics than relying on single data sources alone.

### 3.2. Mortality in black-headed gulls in the wintering areas

Black-headed gulls that had hatched or bred in Germany spent their winters primarily in Central and Western Europe, with wintering areas extending from Ireland in the west to Spain in the south and Poland in the east (**Supplementary Figure S1**). There was a noticeable trend that birds from eastern Germany tended to winter in more eastern regions, ranging from the German Baltic Sea to Croatia and the Adriatic Sea. However, the Western European wintering areas were shared by birds from all German breeding sites, and were located in the Netherlands and Belgium (*N* = 692 ring records, 36.5%), Great Britain and Ireland (*N* = 302, 15.9%), France (*N* = 281, 14.8%), western Germany (*N* = 165, 8.7%), and Spain (*N* = 148, 7.8%).

Black-headed gulls arrive at their wintering areas between August and mid-November and remain there with little individual movement from November to February (Bairlein et al., 2014; Heinicke et al., 2016) (**Figure 2**). By the time black-headed gulls reached these areas in 2022, HPAIV genotype euBB was already circulating there (Briand et al., 2025). In November and December 2022, cases were reported across the entire wintering range of black-headed gulls, although they predominantly affected other species (**Figures 2A, 2B**, **Supplementary Table S2**). In January 2023, the virus still affected other species, but also appeared to have entered (black-headed) gull populations on several independent occasions, with confirmed cases reported in France (*N* = 41), Belgium (*N* = 23), the Netherlands (*N* = 3), Austria (*N* = 3), Spain (*N* = 2), Switzerland (*N* = 2), the United Kingdom (*N* = 2), and the Republic of Ireland (*N* = 1). Outbreaks in black-headed gulls were spatially clustered with those in other species (**Figure 2C**), suggesting interspecies transmission. By February 2023, cases in black-headed gulls had quadrupled, while cases in other species remained stable (**Supplementary Table S2**). Black-headed gulls had thus become the major host species of HPAIV genotype euBB, with major outbreaks now reported in France (*N* = 141), Belgium (*N* = 56), the Netherlands (*N* = 45), Switzerland (*N* = 29), and Italy (*N* = 43) (**Figure 2D**). To conclude, once the virus entered overwintering black-headed gull populations, it spread rapidly with considerably increased mortality.

**Figure 2:**
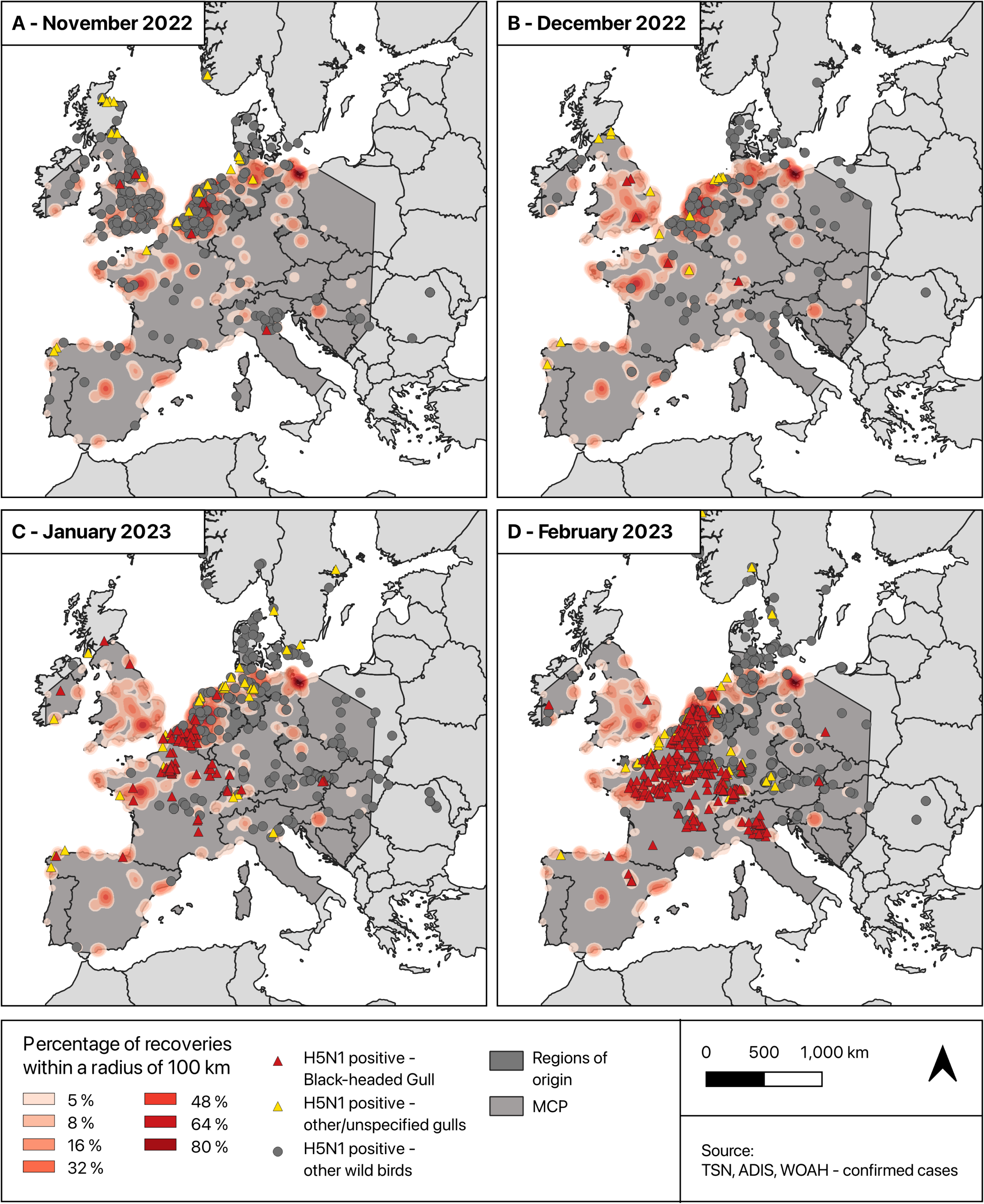
Wintering areas of black-headed gulls of the German breeding population as determined from ring recovery data (displayed in red shades), with confirmed HPAIV outbreak locations overlaid as red triangles (black-headed gulls), yellow triangles (other and unspecified gull species, possibly including also black-headed gulls), and dark grey dots (non-gull species). The winter distribution of black-headed gulls remains constant from November through February, and is thus depicted as stationary. The dark grey polygon represents the minimum convex polygon (MCP) encompassing all winter recoveries of black-headed gulls.

### 3.3. Spatial distribution and timeline of HPAIV-related mortality in black-headed gulls and common terns breeding in Germany in 2023

#### 3.3.1. Impact and spatiotemporal spread of HPAIV incursions into colonies

In Germany, black-headed gulls and common terns often co-occur and share the same breeding habitats, with their primary breeding grounds along the coasts of the North and Baltic Sea (Gedeon et al., 2014). They are also found far inland, particularly across eastern Germany, and extending south to the foothills of the Alps (**Figures 3A, 3D**). In 2023, HPAIV-related mortality was reported across the entire German distribution range of both species. A colony was classified as HPAIV-affected if adults or juveniles either tested positive for HPAIV (*N* = 42 black-headed gull and *N* = 34 common tern colonies), exhibited unusual mortality, or displayed neurological clinical signs suggestive of HPAIV infection, such as debilitation, lethargy, disorientation, loss of flight, opisthotonos, torticollis, and other abnormal behaviours, which typically lead to death within hours of the onset of clinical symptoms (*N* = 10 black-headed gull and *N* = 14 common tern colonies).

**Figure 3:**
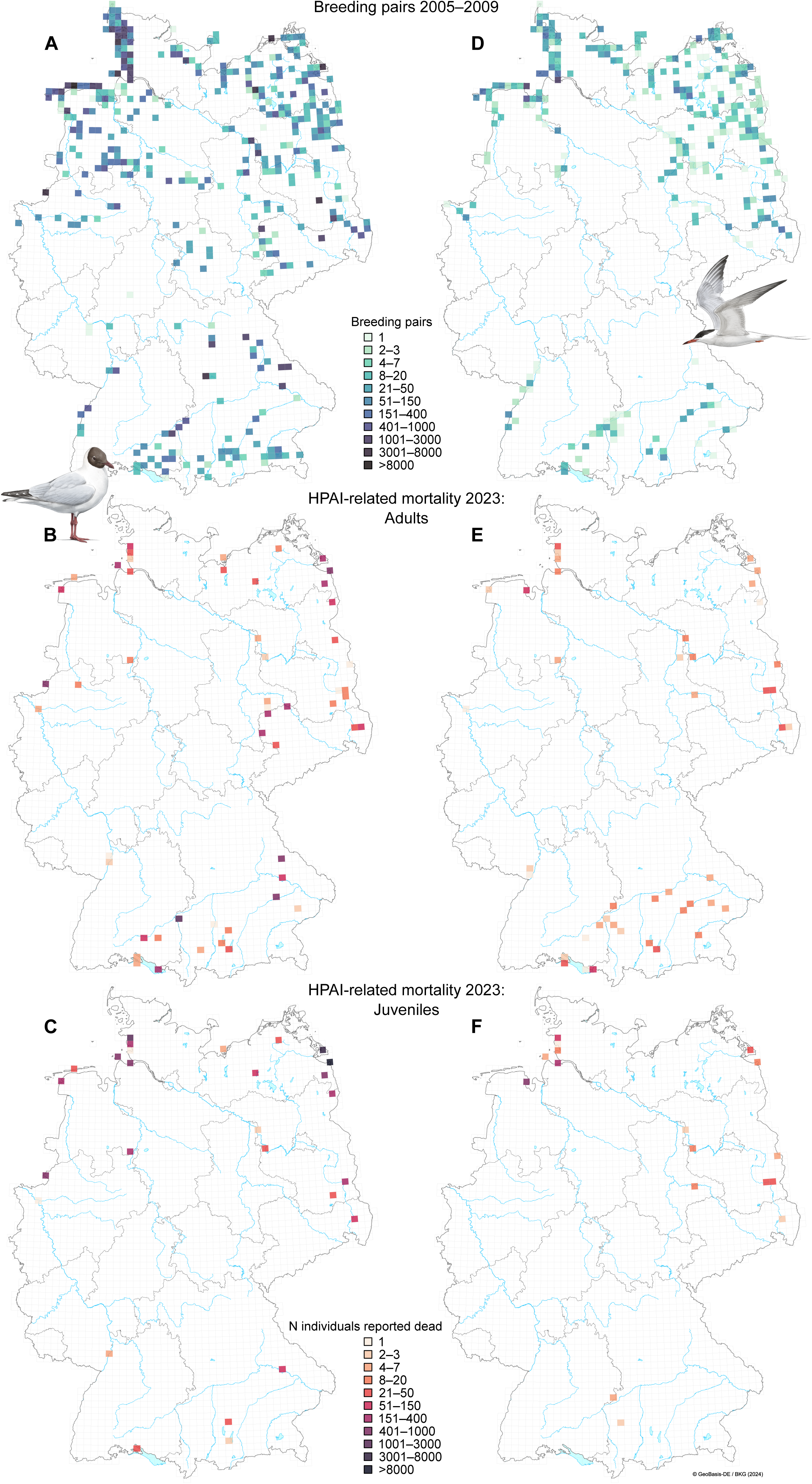
Distribution of black-headed gull and common tern breeding colonies and HPAIV outbreaks in Germany. (A, D) Locations and semiquantitative sizes of breeding colonies of black-headed gulls and common terns, respectively, during 2005–2009 within the German topographical map 1:25,000 raster (TK25, data from Gedeon et al., 2014). (B, E) Semiquantitative HPAI-related mortality among adults in 2023 within the same raster. (C, F) Semiquantitative HPAI-related mortality among chicks in 2023 within the same raster.

Thus, in total, at least 52 colonies of black-headed gulls and 48 colonies of common terns were affected, with 29 colonies shared between the two species (40.8% of all affected colonies). This proportion of colonies where both species co-occurred among HPAIV-affected colonies in 2023 did not differ from the proportion observed in all surveyed colonies, regardless of HPAIV presence, in 2022 and 2023 (Fisher’s exact test: *P* = 0.89 in 2022, *P* = 0.66 in 2023). This suggests that the risk of HPAIV affecting a common tern colony was not elevated by the presence of a black-headed gull colony at the same location (or vice versa).

In total, 8,137 adult black-headed gulls and 614 adult common terns were reported as having been found dead (**Figures 3B, 3E**), along with 29,812 black-headed gull chicks and 1,166 common tern chicks (**Figures 3C, 3F**). Assuming that breeding pair numbers from the most recent national census (conducted approximately 10 years ago; Ryslavy et al., 2020) remain valid, these mortality figures correspond to 3.0% (mean; range: 2.5–3.5%) of the breeding black-headed gulls and 3.5% (mean; range: 3.4–3.6%) of the breeding common terns. Common tern and possibly also black-headed gull populations, however, seem to have declined over the past decade (Südbeck & Packmor, 2024), suggesting that these percentages may underestimate the true impact of HPAIV-related mortality (see also below). In addition, it is important to emphasize that not every bird that died of HPAIV might have been found and reported, such that this percentage represents a minimum estimate of mortality, and the actual value is likely to be considerably higher (Klaassen & Wille, 2023; Knief et al., 2025).

Adult birds were affected across Germany, whereas chick mortality was predominantly observed in the north (**Figures 3B, 3C, 3E, 3F**). This pattern likely resulted from the temporal spread of HPAIV across Germany. The infection wave apparently moved from south to north and from west to east (see also the paragraph on viral phylogenetics below). It initially affected black-headed gulls and started to impact colonies in the south and west by mid-to late-April (**Figures 4A, 4B**). That time of the year typically coincides with the onset of egg laying activity in black-headed gulls (Glutz von Blotzheim, 1999a). Within colonies, infection waves took around one to two weeks to decimate a major fraction of the breeding adults (**Figure 4B**), such that chicks could not have been affected simply because they had not yet hatched. Across colonies, the infection wave progressed north and eastwards, reaching northern Germany within a month. By that time, chicks in the northern colonies had already hatched (Köhler & Neubauer, 2015) and were thus susceptible to either HPAIV infection or starvation due to mortality among their parents.

**Figure 4:**
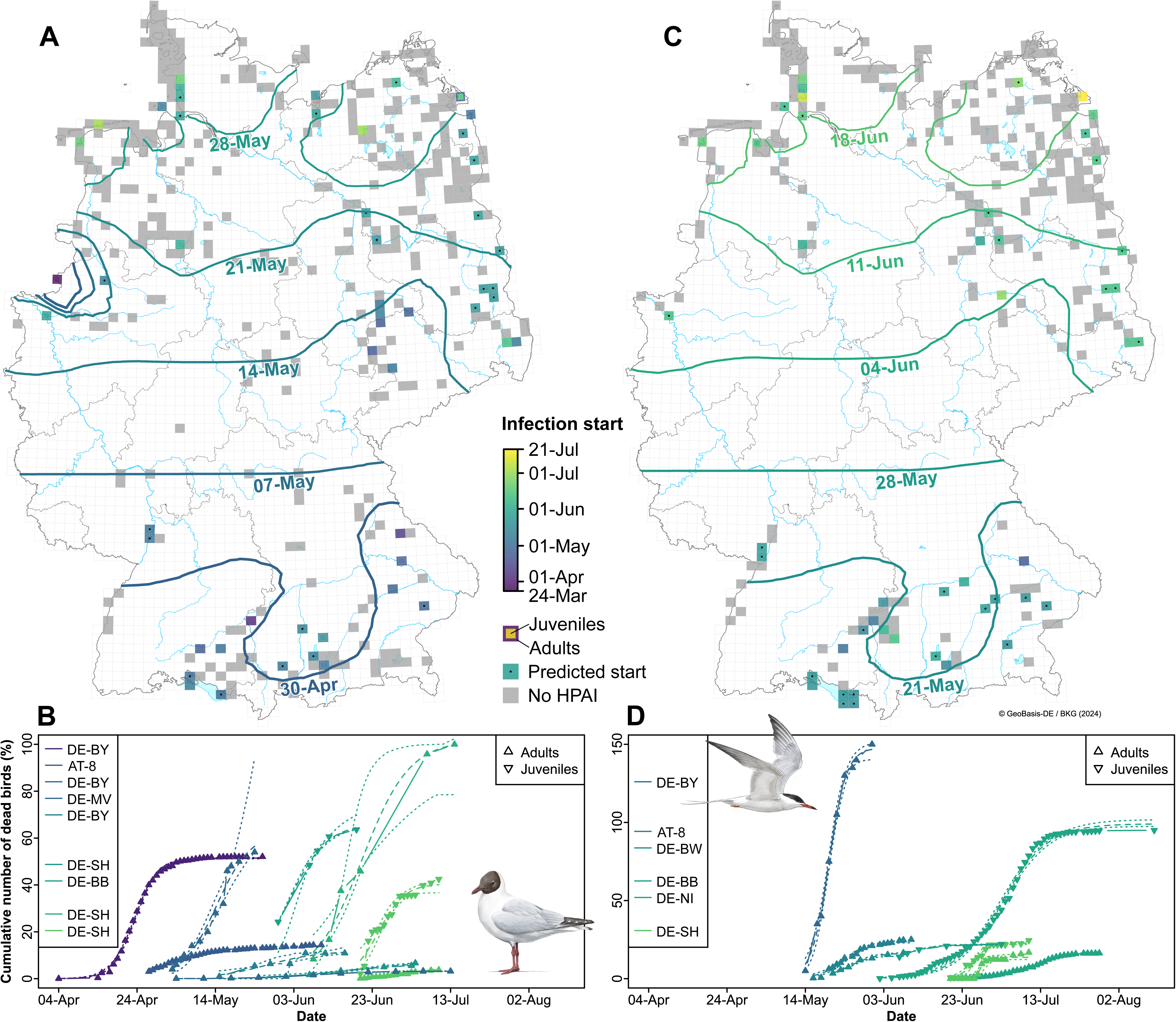
Spatial and temporal dynamics of HPAIV outbreaks across Germany in 2023. (A, C) HPAIV spatial velocity in black-headed gulls and common terns, respectively, depicting the dates of first fatalities within each TK25 raster cell. Raster cells affected by HPAIV, where the dates of first fatalities were missing but predicted using a spatially explicit mixed-effects model, are marked with a dot. Raster cells showing different dates of first fatalities between adults and chicks are two-coloured. Raster cells where black-headed gulls or common terns bred during 2005– 2009, but were not affected by HPAIV in 2023, are shown in grey. Equal-day contour lines illustrate the spatial and temporal spread of HPAIV outbreaks (slightly modified for illustrative purposes here). (B, D) Trajectories of HPAI outbreak waves in specific black-headed gull and common tern colonies, respectively, with data separated into adults and juveniles. Line colours correspond to the dates of first fatalities. Each line represents a single colony and legend entries refer to locations (official two letter country and federal state codes): DE: Germany, AT: Austria, BY: Bavaria, MV: Mecklenburg-Western Pomerania, SH: Schleswig-Holstein, BB: Brandenburg, BW: Baden-Württemberg, NI: Lower Saxony. 8: Vorarlberg. Entries in the legend of (C) aligning horizontally with entries in the legend of (D) refer to the same location /colony.

Black-headed gulls usually arrive at the breeding grounds about a month earlier than long-distance migratory common terns (Glutz von Blotzheim, 1999a, 1999b). Correspondingly, common terns were affected approximately three weeks after the black-headed gulls (spatially-explicit LMM, β ± SE = 20.8 ± 3.8 days, *P* = 6 × 10^−8^, **Figures 4C, 4D**). While infections also appeared to progress from south to north, there was no apparent influx from the west. Despite the later arrival of common terns at the breeding grounds, incubation and nestling periods of gulls and terns in mixed colonies are partially overlapping (Glutz von Blotzheim, 1999a, 1999b). Consequently, similar to black-headed gulls, chick mortality in common terns was only observed in the north, as southern colonies had already collapsed by the time chicks would have begun to hatch.

Given the timing of infections in black-headed gulls and common terns, it appeared that the virus entered the breeding colonies via black-headed gulls and subsequently spilled over to common terns either through direct contact or environmental contamination. Both routes seem plausible. Black-headed gulls prey on common tern eggs (Indykiewicz & Minias, 2019; Robinson, 1999), provoking aggressive defensive behaviours that involve direct contact. Additionally, they parasitize food that common terns bring to their chicks (Brockmann & Barnard, 1979). Moreover, HPAIV-contaminated faeces from black-headed gulls were likely present in the breeding environment of common terns, as both species share the same roosting sites. While these transmission pathways may play a role within colonies, other factors likely determined the risk to which common tern colonies were ultimately affected (see above).

#### 3.3.2. Impact and temporal spread of HPAIV within affected colonies

Within HPAIV-affected colonies, infections were fatal to a substantial fraction of breeding adults, with an estimated 14.8% (negative binomial GLMM, 95% confidence interval: 8.4%–26.2%) of black-headed gulls and 23.0% (negative binomial GLMM, 95% confidence interval: 12.6%–41.9%) of common terns reported dead per colony (see **Figures 4B, 4D**). While this mortality fraction (also called magnitude of outbreaks in the following) was higher in common terns than in black-headed gulls, the difference was not statistically significant (*P* = 0.19, excluding seasonal effects), and mostly driven by black-headed gulls – but not common terns (*P* = 0.11) – having higher mortality rates at the onset of the season (negative binomial GLMM, each additional day was associated with a multiplicative change in the expected number of dead birds of 0.98 (95% CI: 0.96–1.00), corresponding to a 2.0% daily decline, *P* = 0.048).

Colony identity explained a significant proportion of the variance in this mortality fraction (*P* = 0.034), indicating that infection severity was correlated between species: when black-headed gulls were heavily affected in a colony, common terns were also heavily affected, and vice versa (intraclass correlation coefficient = 0.32). This suggests that the environmental viral load affects both species similarly.

In the 12 colonies for which we had within-season longitudinal data (**Figures 4B, 4D**), we found that fatality rates (which are the slopes in **Figures 4B, 4D**) were higher in common terns than in black-headed gulls (LMM, β ± SE = 0.14 ± 0.04, *P* = 0.022). In this subset of colonies, there was no evidence that either the magnitude of an outbreak or the fatality rate changed with seasonal progression (both *P* > 0.3) or with breeding pair density (both *P* > 0.06).

To conclude, common terns appeared to be more severely affected than black-headed gulls, both in terms of mortality fraction (i.e., the asymptotes in **Figures 4B, 4D**) and fatality rates (i.e., the slopes in **Figures 4B, 4D**), with some indication of seasonal attenuation and colony-specific effects.

### 3.4. Phylogenetic and spatiotemporal analyses of the HPAIV genotype euBB in gull and tern species

The gull-adapted HPAIV genotype euBB emerged in summer 2022 in France and spread throughout Europe; it was first detected in Germany in 2023. We sequenced 25 German genotype euBB samples and aligned them with the hemagglutinin (HA) coding sequences from 264 European H5N1 genotype euBB viruses, collected from various gull and tern species, to build a dataset of 289 HA sequences. This was used to construct a time-scaled maximum clade credibility (MCC) phylogeny and to conduct a phylogeographic analysis estimating the virus’ spread across Europe, with a particular emphasis on its spatiotemporal dynamics in Germany. The MCC phylogeny indicated that German viral genomes shared a common ancestor dating back to mid-2022 (**Supplementary Figure S2**), suggesting the virus diversified outside of Germany. Interestingly, samples from northern German federal states (Schleswig-Holstein, Lower Saxony, Mecklenburg-Western Pomerania) formed a distinct branch suggesting distinct entry and spreading pathways of genotype euBB variants. This pattern is also partially reflected in the bird ringing and recovery data (**Supplementary Figure S1**).

Spatiotemporal phylogeographic analysis of viruses sampled between November 2022 and December 2023 revealed seasonal patterns of a southward spread during winter (indicated by arrows in shades of blue in **Figure 5**) and northward spread during summer months (arrows in shades of red). Western Germany was affected by incursions from the west, originating from the major wintering grounds of black-headed gulls in the Netherlands and Belgium (**Figure 5**). Southern and eastern Germany, however, were reached predominantly from Poland and Switzerland (**Figure 5**). The spatiotemporally-resolved analysis confirmed several independent incursions into Germany, consistent with the time-scaled phylogeny. The first wave began in March and April 2023, with HPAIV spreading from Poland to Schleswig-Holstein and Mecklenburg-Western Pomerania, covering the eastern and northern federal states of Germany, and extending to Baden-Württemberg in the south. An additional incursion from Switzerland into Bavaria was observed in April. In western Germany, North Rhine-Westphalia experienced introductions from the Netherlands during March and April. A second wave followed in May and June 2023, again showing eastward spread from the Netherlands into Lower Saxony and Baden-Württemberg, and westward movements from Poland into Mecklenburg-Western Pomerania, Saxony, and Saxony-Anhalt. These patterns broadly align with the observed spatiotemporal distribution of HPAIV infections in German black-headed gulls and common tern colonies; however, time-resolved mortality data from neighbouring countries are lacking to support the inferred spatiotemporal spread pattern based on both viral and mortality data.

**Figure 5:**
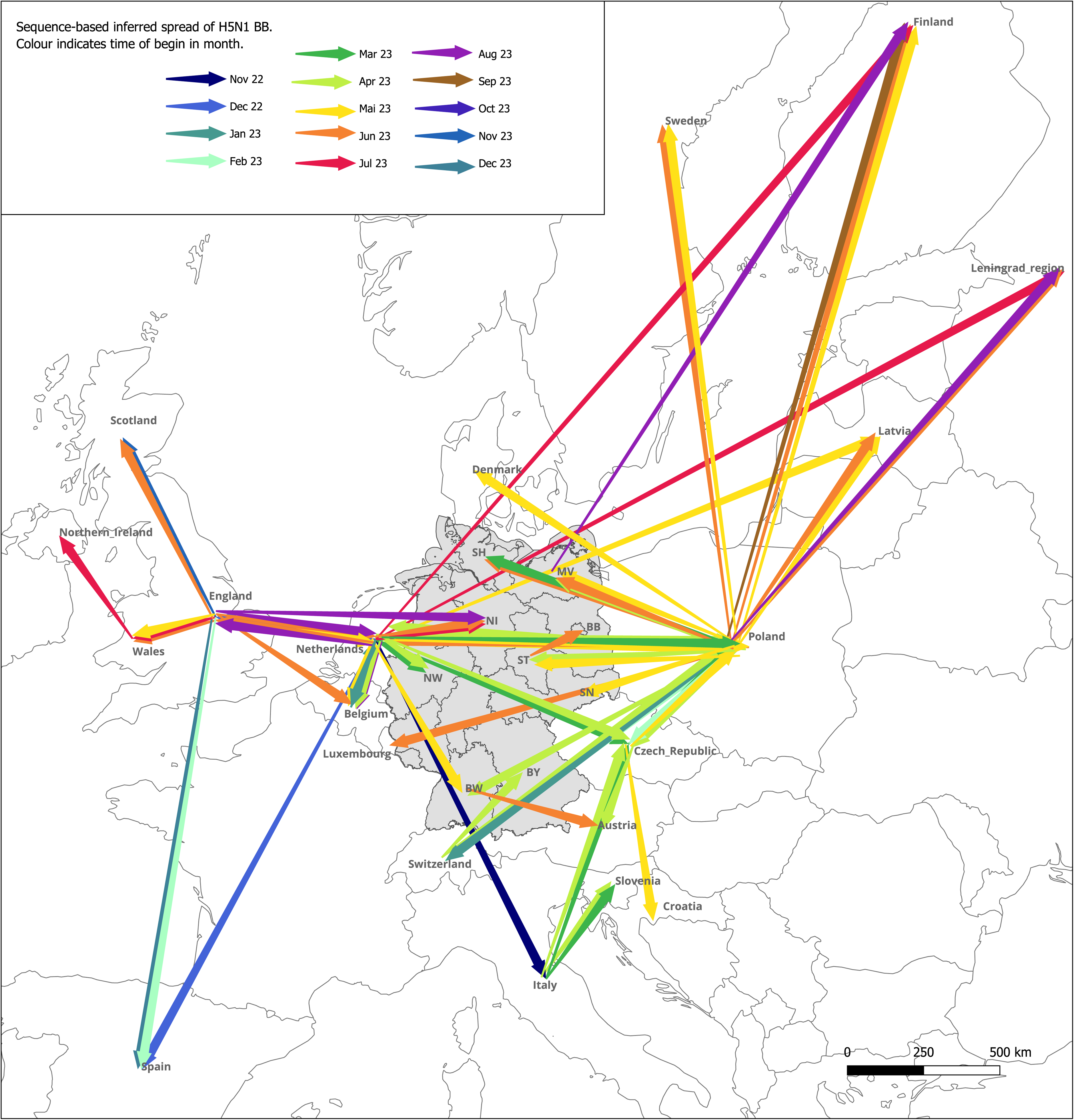
Spatiotemporal spread of the HPAIV H5N1 genotype euBB across Europe and incursion into Germany (greyed area) between November 2022 and December 2023, inferred by HA gene sequence analysis. Arrows indicate inferred viral incursions into, and spread within, European countries and German federal states, but do not represent movements of individual birds or avian populations. Arrow colours distinguish seasons (shades of blue: winter, shades of green: spring, shades of red: summer). German federal states are shown with abbreviated names (SH: Schleswig-Holstein, NI: Lower Saxony, MV: Mecklenburg-Western Pomerania, ST: Saxony-Anhalt, SN: Saxony, BB: Brandenburg, NW: North Rhine-Westphalia, BW: Baden-Württemberg, BY: Bavaria).

### 3.5. Active surveillance for HPAIV in breeding colonies of black-headed gulls and common terns in northern Germany

To assess the number of individuals surviving HPAIV H5 exposure, antibodies against HPAIV (H5) were measured in serological surveys conducted in breeding colonies of black-headed gulls and common terns at two sites in northeastern and northwestern Germany. These surveys aimed to detect both general and H5-specific antibodies in potentially affected colonies, providing insights into past infections and the presence of potentially protective immunity within these populations.

In 2023, during the HPAIV outbreak, we collected swab and serum samples from 25 apparently healthy adult black-headed gulls in the breeding colony on the Isle of Böhmke in northeastern Germany. Additionally, we obtained 739 serum samples from apparently healthy adult (*N* = 573) and 20-day old chick (*N* = 155) common terns at the Banter See colony in northwestern Germany, where sampling was conducted during the HPAIV-induced mass mortality events in 2022 and 2023, as well as a year later when no birds died from an HPAIV infection (**Figure 6; Supplementary Table S3**). We screened all serum samples for the presence of antibodies, and also tested black-headed gulls for influenza A virus (IAV)-specific RNA to detect active infections.

**Figure 6:**
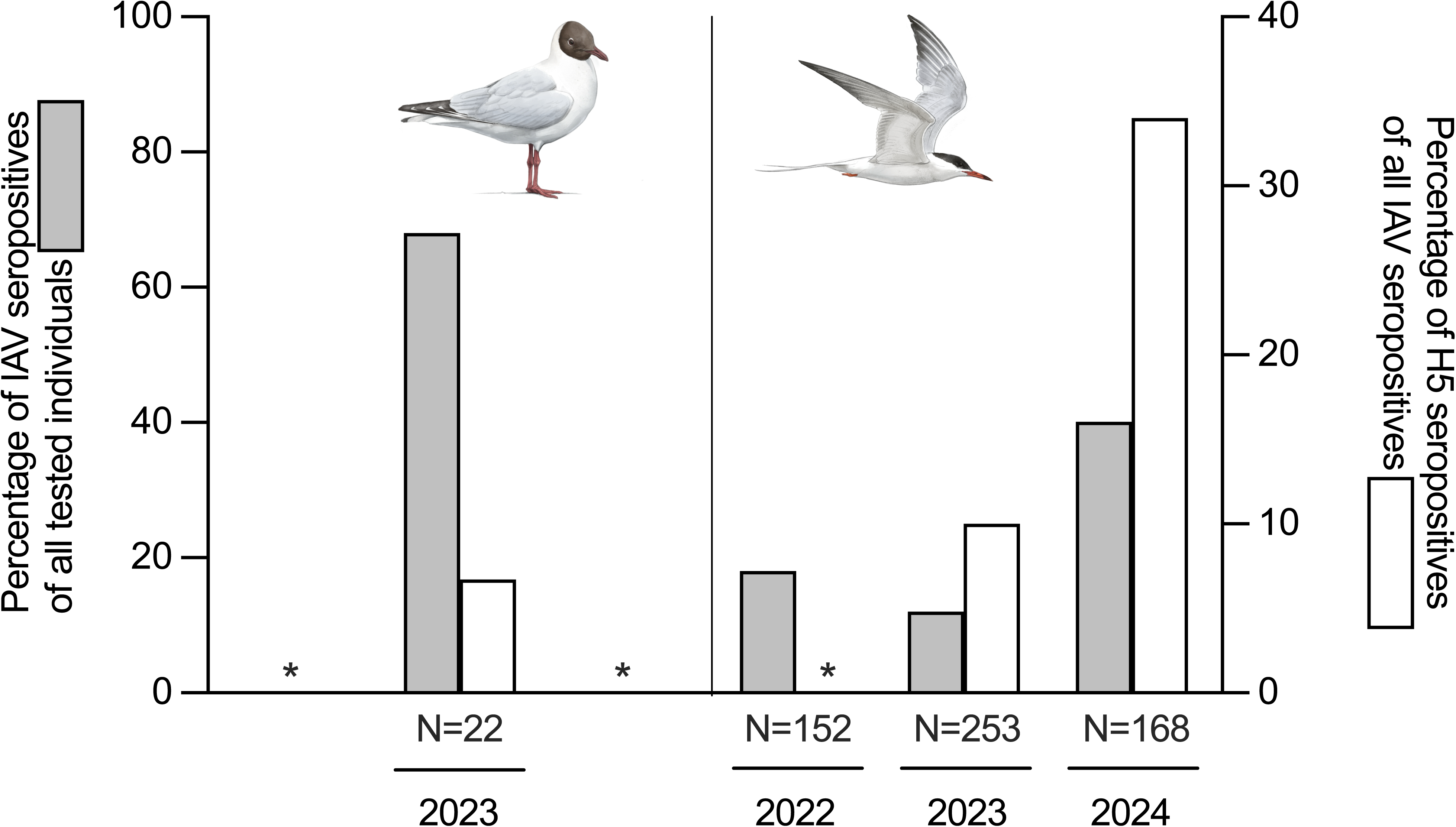
Seroreactivity to influenza A virus (IAV) and H5 subtype-specific antibodies in adult breeding black-headed gulls (left panel) and common terns (right panel) sampled between 2022 and 2024. Grey bars indicate the percentage of IAV-seropositive individuals among all tested individuals (left Y-axis), while white bars show the percentage of H5-seropositive individuals among IAV-seropositives (right Y-axis). * = No samples available for testing.

The majority of black-headed gulls tested negative for IAV-specific RNA in generic RT-qPCR screenings. Only two samples showed a positive signal with high Ct-values (37.7 and 38.3), indicating low viral loads for an avian influenza virus. In subsequent RT-qPCR tests, subtypes H5, H13, H16 and N1 were excluded, confirming that these birds were not infected with HPAIV H5N1. However, one dead black-headed gull found at the same sampling location tested positive for HPAIV H5N1, with Ct-values ranging from 25.2 to 27.1 in different tissues. IAV-specific antibodies were detected in 68% of the black-headed gulls (*N* = 15 out of 22 serum-sampled individuals), but H5-specific antibodies were confirmed by ELISA in only one individual (**Figure 6**; **Supplementary Table S3**).

Among breeding adult common terns, 18% and 12% had antibodies against the IAV-specific NP protein in 2022 and 2023, respectively. This proportion increased to 40% in 2024. This rise in NP-specific seropositive birds was accompanied by a gross increase of birds that tested positive for H5-specific antibodies (**Figure 6**). Of the 155 chicks sampled in 2023 and 2024, only two had IAV-specific antibodies, but both scored negative for H5-specific antibodies.

Results showed that, particularly in common terns, HPAIV H5N1 infection is not invariably lethal and that a substantial proportion of the adult common terns at the Banter See developed a humoral H5-specific immune response.

### 3.6. Population impact of HPAIV on black-headed gull and common tern survival and breeding pair numbers

HPAIV-related mortality in 2023 was substantially higher than the background mortality in years without HPAIV. Estimated apparent annual survival rates for adults of both species prior to 2022 are around 0.85–0.90 (range in black-headed gulls: 0.73– 0.90, in common terns: 0.85–0.95 (Breton et al., 2014; DiCostanzo, 1980; Heldbjerg, 2001; Nisbet & Cam, 2002; Ottersland, 2024; Palestis & Hines, 2015; Prévot-Julliard et al., 1998; Szostek & Becker, 2012), whereas we found an additional mortality rate of 0.030 and 0.035 within the c. five months of breeding season in 2023 (see above and **Supplementary Figure S4**). This suggests that mortality was at least 30-35% higher in 2023 than in years without HPAIV-related mass mortality. To illustrate this, we calculated the average monthly numbers of black-headed gulls and common terns ringed and reported dead in eastern Germany from 2010 to 2022 and compared them to those reported dead in 2023. Mortality was considerably higher throughout the breeding season (March to July) in 2023 compared to previous years (**Supplementary Figure S4**).

To estimate the impact of HPAIV-related mass mortality on the number of breeding pairs in the following year, we compiled datasets of breeding pair (BP) numbers of black-headed gulls and common terns in Germany between 2022 and 2024. These data were not complete, but captured around 55.5–77.2% of the black-headed gull population and 83.2–88.1% of the common tern population based on the German 2012–2016 breeding pair estimates (Ryslavy et al., 2020). However, given the completeness of our common tern dataset, it is likely that nearly all breeding pairs were included, and that the observed decline of more than 10% since 2016 reflects a genuine population reduction. Suitable breeding habitat was created in the Szczecin Lagoon in Poland in recent years, attracting substantial numbers of both black-headed gulls and common terns (Marchowski et al., 2024). These newly formed colonies may have led to emigration from the eastern parts of Mecklenburg-Western Pomerania, potentially blurring the apparent population impact of HPAIV on breeding pair numbers. We therefore excluded data from eastern Mecklenburg-Western Pomerania from the following calculations, unless stated otherwise (see **Supplementary Figure S3** for an illustration of the full dataset). It is worth noting, however, that both species declined substantially in this region of Germany: in 2022, black-headed gulls numbered 22,638 BP and common terns 516 BP; in 2023, 22,589 BP and 382 BP, respectively; and in 2024, 11,849 BP and 153 BP. In black-headed gulls, the loss in eastern Mecklenburg-Western Pomerania (−10,789 BP) could not be offset by the increase in the Polish Szczecin Lagoon (+3,219 BP), and was comparable to the decline across the rest of Mecklenburg-Western Pomerania (−47.7% vs −47.0% in BP numbers between 2022 and 2024, in the east and the rest of the region, respectively). In contrast, the decline in common terns (−363 BP) was more than compensated by the increase in Polish breeding pairs in the Szczecin Lagoon (+823 BP) and much stronger than in the rest of Mecklenburg-Western Pomerania (−70.3% vs −17.6% in BP numbers between 2022 and 2024, in the east and the rest of the region, respectively).

Across Germany, black-headed gull BP numbers declined by 15.9% between 2022 and 2024 (from 66,139 to 55,622; *P* = 0.028), while common terns declined by 5.8% over the same period (from 6,972 to 6,569; *P* = 0.060; **Figures 7A, 7B**). In black-headed gulls, inland colonies experienced a significant decline (estimated log difference between 2022 and 2024 = −0.44, SE = 0.14, *P* = 0.013), whereas no significant change was detected in coastal colonies overall. However, the raw data reveal a substantial decline in coastal Mecklenburg-Western Pomerania (see above), indicating that regional effects may be masked when aggregating coastal data. In common terns, inland colonies also showed a significant decline between 2022 and 2024 (estimated log difference = −0.43, SE = 0.12, *P* = 5.1 × 10^−3^), whereas coastal colonies remained relatively stable. Yet, the coastal colony at the Banter See in Lower Saxony illustrates the severe impact of HPAIV: breeding pair numbers dropped from 690 in 2022 to 340 in 2023 and 215 in 2024, following outbreaks in both 2022 and 2023.

**Figure 7:**
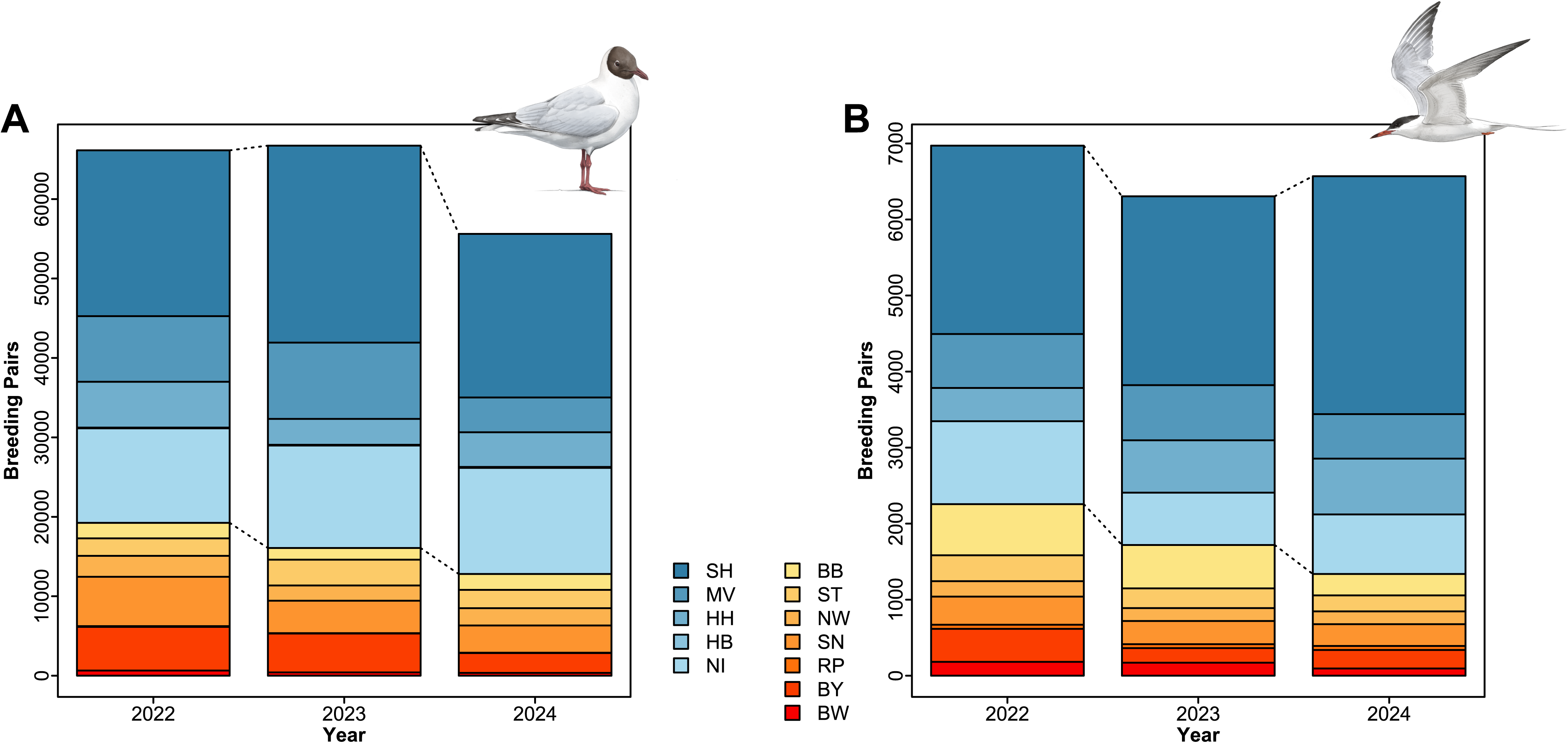
Changes in breeding pair numbers of (A) black-headed gulls and (B) common terns per federal state in Germany from 2022 to 2024. Data from the eastern part of Mecklenburg-Western Pomerania were excluded due to substantial cross-border movements to newly established colonies in the Szczecin Lagoon, Poland (see Supplementary Figure S3 for the full dataset). Colours distinguish inland colonies (shades of red to yellow) from coastal ones (shades of blue). Note that the figure does not represent the entire German breeding population; data from inland Lower Saxony, and several other regions are missing. Relative to the most recent national census, our 2022 dataset represents 55.5–77.2% of the total number of breeding pairs in black-headed gulls and 83.2–88.1% in common terns. German federal states are shown with abbreviated names (SH: Schleswig-Holstein, MV: Mecklenburg-Western Pomerania, HH: Hamburg, HB: Bremen, NI: Lower Saxony, BB: Brandenburg, ST: Saxony-Anhalt, NW: North Rhine-Westphalia, SN: Saxony, RP: Rhineland-Palatinate, BY: Bavaria, BW: Baden-Württemberg).

Together, these patterns suggest that coastal colonies of both species may have been less affected, potentially due to milder outbreaks along the coasts in 2023 or redistribution of birds from inland to coastal colonies where breeding conditions may be more favorable. A similar trend has been reported for Sandwich terns, where colonies in the most severely affected areas experienced smaller declines than adjacent sites.

## 4. Discussion and Conclusions

We report on the epidemiology of the HPAIV H5N1 2.3.4.4b genotype euBB in black-headed gulls and common terns in Germany, which began in the winter of 2022/23 and continued throughout the 2023 breeding season. Genotype euBB emerged from a reassortment event involving a HPAIV H5N1 and a gull-adapted LPAIV in northern France in the spring of 2022 (Briand et al., 2025; Fusaro et al., 2024), and quickly dominated the epidemiological landscape on the wintering grounds of German black-headed gulls in the Netherlands, Belgium and France in the winter of 2022/23.

We show that black-headed gulls rapidly became an important host species and most likely introduced the virus to the German breeding areas through multiple incursions from the Western European wintering grounds. Other waterbirds like anseriform species and scavenging raptors may have also contributed to the spread (Caliendo et al., 2025; van den Brand et al., 2015) (see also **Supplementary Figure S5**). This means that there is no “gull-specific” viral phylogenetic lineage that would suggest that it only spreads within black-headed gulls and from colony to colony. Subsequently, the virus spilled over to common terns, which frequently share breeding colonies with black-headed gulls and have a similar breeding phenology, but typically arrive at the colonies about three to four weeks later (Glutz von Blotzheim, 1999a, 1999b), which is consistent with the approximately 21-days delay in the onset of mass mortality observed in this species (**Figure 4**).

This later arrival of common terns is explained by differences in life history between the two species. As we have shown, German black-headed gulls are partial short-distance migrants that only rarely reach North and West Africa (Bairlein et al., 2014), and they return to their breeding colonies between March and mid-April (Glutz von Blotzheim, 1999a). In contrast, common terns are long-distance migrants. Recent geolocator studies revealed two distinct migration routes: most common terns breeding in Germany follow the Atlantic coast to West Africa (95%) or southern Africa (5%) (Kürten et al., 2022), but a small proportion of terns from eastern Germany takes a different route, via the eastern Mediterranean Sea, the Red Sea and East African coasts to Southeast or southern Africa (Piro & Schmitz Ornés, 2022). After migration, common terns return to their breeding colonies later than black-headed gulls, typically during the second half of April or early May (Glutz von Blotzheim, 1999b). Consequently, common terns lay and hatch their eggs later than black-headed gulls, but due to incubation and chick rearing lasting 20–26 days and 23–28 days, respectively (Glutz von Blotzheim, 1999a, 1999b), both species show a strong phenological overlap at the breeding colonies. Birds of both species leave their breeding colonies from mid-July. Common terns often begin their autumn migration shortly afterwards, but some remain at favourable (stopover) sites in Central Europe until early September.

The spatial and temporal spread of HPAIV broadly aligns with the migratory movements of black-headed gulls. In both the 2022/23 and 2023/24 autumn and winter seasons, the virus moved southward to Spain and Italy. It returned from Italy to Central Europe in spring 2023, but subsequently spread further north during the summer, a period when European black-headed gulls are largely stationary and do not migrate (Bairlein et al., 2014). Several factors may account for this pattern. First, the virus was not restricted to black-headed gulls, but also infected other species that may have facilitated its continued northward spread. Second, studies using GPS-tagged seabirds have shown that some individuals surviving an HPAIV outbreak dispersed over large distances (Careen et al., 2024; Jeglinski et al., 2024). If similar post-outbreak movements occurred in black-headed gulls, they may have contributed to the northward spread of the virus later in the season. Third, although breeding black-headed gulls typically do not move over large distances (Jakubas et al., 2020), non-breeding adults, individuals whose breeding attempts have failed, and young birds may engage in prospecting behaviour in search of suitable colonies, potentially covering large geographic areas and facilitating further viral dissemination beyond the typical migration season (Boulinier, 2023; Oro et al., 2021). Alternatively, the reconstructed timing of viral spread may reflect delayed detection. The virus could have arrived during the regular migration period in northern Europe, but circulated undetected in local populations, or mortality occurred earlier but carcasses were only found and tested one or two months later.

In total, 8,137 adult black-headed gulls (3.0% of the breeding birds) and 614 adult common terns (3.5% of the breeding birds) were reported dead in German breeding colonies in 2023, but this number likely represents only the tip of the iceberg, as the decline in breeding pair numbers in 2024 was substantially greater than expected based on this mortality. The discrepancy could be explained by unrecorded mortality in non-surveyed colonies or outside of the breeding sites, a lack of reporting by site managers, or mortality occurring at the wintering grounds or during migration (Jatta et al., 2025).

What ultimately matters are the actual breeding pair numbers, which declined by 15.9% and 5.8% in black-headed gulls and common terns in Germany between 2022 and 2024. When data from eastern Mecklenburg-Western Pomerania are included, these declines increase to 24.0% and 10.2%, respectively. Assuming that our dataset captured nearly all breeding pairs of common terns in Germany in 2022 and subsequent years, the national population declined by more than 10% between 2016 and 2022, and by over 20% by 2024. These recent declines reflect a combination of an already negative long-term trend, now exacerbated by HPAIV-related mortality. In addition, some birds may have relocated to more suitable breeding regions (e.g., the Szczecin Lagoon in Poland), further complicating the interpretation of national trends. Taken together, these findings suggest that the conservation status of the common tern in the German Red List of endangered avian species (Gerlach et al., 2019; Ryslavy et al., 2020) may warrant reassessment, potentially requiring an upgrade from “Highly Threatened” to “Threatened with Extinction” (for a detailed explanation see **Supplementary Text**). The black-headed gull remains a common breeding bird in Germany, with a previously stable breeding population (Gerlach et al., 2019; Ryslavy et al., 2020). It could remain classified as “Not Threatened”, despite the substantial reduction in breeding pair numbers of 15–24% within just two years.

Recorded mortality was higher among common terns than black-headed gulls, yet the decline in the number of breeding pairs was more pronounced in black-headed gulls. This discrepancy may have two explanations: First, mass mortality in black-headed gulls occurred for the first time in 2023, whereas common terns had already experienced high mortality in coastal colonies in 2022 (Pohlmann et al., 2023), likely due to spillover from other severely affected species such as Sandwich terns, which breed at the same sites (Knief et al., 2024). While this earlier mortality may have already reduced breeding pair numbers in common terns by 2022, available data do not support this interpretation. Second, there may be a reporting bias: as an endangered species in Germany, common tern colonies are monitored more intensively than those of black-headed gulls, potentially leading to higher detection rates of dead individuals. This underscores the importance of monitoring across all affected colonies and species, also during acute HPAIV-induced mass mortality events.

We did not find an association between outbreak size and environmental variables such as breeding bird density, as observed in other countries (Marchowski et al., 2025) and other species (Bregnballe et al., 2024; Lane et al., 2024). The absence of such an effect may be caused by the small sample size and low variation in breeding density in our dataset (mean ± SD density in back-headed gulls = 0.54 ± 0.32 BP/m^2^, *N* = 9 colonies; mean ± SD density in common terns = 1.06 ± 0.60 BP/m^2^, *N* = 6 colonies).

High adult and chick mortality among black-headed gulls and common terns in 2023 was not limited to Germany, but occurred across the entire European breeding range of both species (Knief et al., 2025). Mass-mortality events were reported from the Czech Republic (Nagy et al., 2024), Finland (Kareinen et al., 2024), France (EFSA (European Food Safety Authority) et al., 2023), the Netherlands (Ballmann & Lilipaly, 2024; de Boer, 2024), Poland (Indykiewicz et al., 2025; Marchowski et al., 2025; Przymencki et al., 2024), the Republic of Ireland (Burke et al., 2024), and the United Kingdom (Owens, 2024; Tremlett et al., 2024). For example, Indykiewicz et al. (2025) surveyed 114 black-headed gull colonies across Poland and estimated an HPAIV-related adult mortality rate of 22.2%. When extrapolated to the national breeding population (at least 115,000 pairs according to the Polish national census), this corresponds to approximately 51,000 adult deaths in 2023. Marchowski et al. (2025) investigated the effects of HPAIV in 2022/23 on colonial waterbirds in the Lower Odra Valley. They estimated a mortality rate of 14% for black-headed gulls (2,484 BP) and 12% for common terns (656 BP), and found a positive relationship between bird density and the extent of mortality.

With the end of the 2023 breeding season, mortality in black-headed gulls and common terns declined rapidly, as did the incidence of HPAIV genotype euBB (Fusaro et al., 2024), which was swiftly replaced by other HPAIV H5N1 2.3.4.4b genotypes (Fusaro et al., 2024), accompanied by a shift in host species range towards waterbirds and raptors (EFSA (European Food Safety Authority) et al., 2025; EFSA (European Food Safety Authority) et al., 2024). The reasons for the turnover of genotype euBB remain speculative: initial host (gull) species may have acquired a degree of population immunity, and other genotypes might have attained a higher fitness in non-charadriiform hosts than genotype euBB. Our serological findings support the presence of H5-subtype specific antibodies in black-headed gulls after exposure to H5N1, and a gradual increase in H5-subtype specific seroprevalence over time in common terns, which appears to be primarily attributable to exposure to H5Nx viruses. Based on studies in model organisms such as mallards (*Anas platyrhynchos*) and chickens (*Gallus gallus*), these antibodies are expected to have a virus neutralizing effect and to contribute to protection against infection and clinical disease (Jourdain et al., 2010). Ongoing studies over the next few years, especially of individually identifiable common terns at Banter See, will provide further information on how long these antibodies remain detectable and whether projections of protection at the population scale can be derived from them. The anticipated repeated exposure of this species to HPAIV H5 over its lifespan, combined with its geographically extensive range, may facilitate the stabilization of an H5-seropositive immune status (and protection) through continual boosting (Arnal et al., 2015). However, it is important to note that widespread population immunity also increases selection pressure on the virus to evolve escape mutants via antigenic drift. These viruses may have higher fitness due to their ability to evade host immunity (Latorre-Margalef et al., 2017; Luczo & Spackman, 2024).

Even if immunity has arisen, both black-headed gulls and common terns have witnessed substantial population declines. Given that both species display a slow life history, population recovery may take decades. In addition, seabird populations face various other threats as well (Phillips et al., 2023): climate change is increasing the frequency of egg and nestling loss due to adverse weather and flooding during the breeding season, while drought and intensified fishing activities may reduce food availability. Pollution may further lower survival rates, and changes in agricultural practices and human activity are contributing to increasing densities of mammalian predators. Additionally, tourism is rendering previously suitable breeding habitats unusable. HPAIV has now been added to this growing list of pressures on seabirds, a group that was already the most threatened among all bird taxa globally prior to the emergence of HPAIV (Paleczny et al., 2015). Should further HPAIV variants or other emerging diseases affect them, their future may be bleak.

## Declarations

### Ethics Approval and Consent to Participate

Paragraph 2.6 on “Active surveillance for HPAIV in black-headed gulls and common terns in northern German” comprises declarations on ethical approval for the active surveillance of Black-headed gull and Common tern populations in Germany.

## Consent for Publication

Not applicable.

## Availability of Data and Material

A detailed overview of reported mortalities and of breeding pair estimates are provided as Supplementary data. Geographical geojson vector maps were obtained from http://opendatalab.de/projects/geojson-utilities/ with open data provided by the German Federal Agency for Cartography and Geodesy (https://gdz.bkg.bund.de/). Virus nucleotide sequence data with accession numbers, associated metadata and acknowledgement of the originating and submitting laboratories are available from Zenodo with identifier doi: 10.55876/gis8.250320fm. Associated information and source data of the phylogenetic and spatiotemporal data are available on Zenodo with identifier doi: 10.5281/zenodo.14713066.

## Acknowledgements

Without the data contributed by dedicated fieldworkers and volunteers, this study would not have been possible. A comprehensive acknowledgment to all those who provided input is presented as a supplement in the additional file.

## Authors Contributions

**UK**: Conceptualization, Data curation, Formal analysis, Investigation, Methodology, Project administration, Validation, Visualization, Writing - original draft, Writing - review & editing

**AKA**: Formal analysis, Investigation

**VA**: Investigation

**CJB**: Formal analysis, Methodology, Visualization

**JB**: Investigation

**SB**: Conceptualization, Investigation, Resources, Writing - review & editing

**WF**: Data curation, Writing - review & editing

**AGl**: Conceptualization, Data curation, Formal analysis, Visualization, Writing - original draft, Writing - review & editing

**AGü**: Formal analysis, Investigation, Methodology, Validation, Visualization, Writing - original draft, Writing - review & editing

**CH**: Data curation, Formal analysis, Investigation, Methodology, Writing – review & editing

**SK**: Investigation, Writing - review & editing

**DM**: Data curation, formal analysis, Writing – review & editing

**SP**: Data curation, Investigation, Methodology, Writing – review & editing

**AP**: Data curation, Formal analysis, Visualization

**RER**: Investigation

**CS**: Data curation, Formal analysis, Investigation, Methodology, Writing – review & editing

**TH**: Conceptualization, Project administration, Supervision, Visualization, Writing - original draft, Writing - review & editing

All authors read and approved the final manuscript.

## Additional Material

### Additional file

“Supplementary Material on *Outbreak dynamics of high pathogenicity avian influenza virus H5N1, clade 2.3.4.4b euBB, in black-headed gulls and common terns in Germany in 2023*”

The additional file provides supplementary text, figures and tables supporting the main manuscript.

