## Supplementary Material for "Outbreak dynamics of high pathogenicity avian influenza virus H5N1, clade 2.3.4.4b euBB, in black-headed gulls and common terns in Germany in 2023"

Ulrich Knief[1,*], Ann Kathrin Ahrens[2], Valerie Allendorf[2], Carla J. Behringer[3], Justine Bertram[4], Sandra Bouwhuis[4], Wolfgang Fiedler[5], Anja Globig[2], Anne Günther[2], Christof Herrmann[6], Sascha Knauf[2,7], Dominik Marchowski[8], Simon Piro[6], Anne Pohlmann[2], Robert E. Rollins[4], Christoph Staubach[2], Timm Harder[2,#]

[1] University of Freiburg, Institute of Biology I (Zoology), Evolutionary Biology and Ecology, Hauptstr. 1, DE-79104 Freiburg, Germany

[2] Friedrich-Loeffler-Institut, Federal Research Institute for Animal Health, Südufer 10, DE-17493 Greifswald-Insel Riems, Germany

[3] Universität Innsbruck, Fakultät für Biologie, Innrain 52, 6020 Innsbruck, Austria

[4] Institute of Avian Research, An der Vogelwarte 21, DE-26386 Wilhelmshaven, Germany

[5] Max Planck Institute of Animal Behavior, Am Obstberg 1, DE-78315 Radolfzell, Germany

[6] Agency for Environment, Nature Conservation, and Geology Mecklenburg-Western Pomerania, Hiddensee Bird Ringing Scheme, Goldberger Str. 12b, DE-18273 Güstrow, Germany

[7] Justus Liebig University, Faculty of Veterinary Medicine, Professorship for One Health/International Animal Health, Frankfurterstr. 106, DE-35393 Giessen, Germany

[8] Ornithological Station, Museum and Institute of Zoology, Polish Academy of Sciences, Nadwiślańska 108, 80-680 Gdańsk, Poland

* Address for correspondence: Ulrich Knief, University of Freiburg, Institute of Biology I (Zoology), Evolutionary Biology and Ecology, Hauptstr. 1, DE-79104 Freiburg, Germany, Phone: 0049-761-203-2911,

[#] Address for correspondence: Timm Harder, Friedrich-Loeffler-Institut, Federal Research Institute for Animal Health, Südufer 10, DE-17493 Greifswald-Insel Riems, Germany, Phone: 0049-38351-7-1546,

### Index


Supplementary text 3

Supplementary figures 4

Supplementary tables 9

Supplementary references 12

Supplementary acknowledgments 13

ORCID references of all authors 15

### Supplementary text

**Population impact of HPAIV on black-headed gull and common tern breeding pair numbers including all German data**

Using the entire German breeding pair numbers, including eastern Mecklenburg-Western Pomerania but excluding the Polish part of the Szczecin Lagoon, black-headed gull breeding pair (BP) numbers declined by 24.0% (from 88,777 in 2022 to 67,471 in 2024; $P = 0.025$), while common terns declined by 10.2% (from 7,488 to 6,722; $P = 0.037$; blue, yellow and red segments of the stacked bars in **Figures S3A**, **S3B**). In the Polish part of the Szczecin Lagoon (Odra River Estuary), two artificial islands have been recently created, offering suitable breeding grounds for black-headed gulls and common terns (Marchowski et al., 2024). These sites have triggered rapid population growth and attracted birds also from other regions of Poland and Germany, likely contributing to the observed increase in common tern numbers between 2023 and 2024, and possibly obscuring the general population trend (see the green segments of the stacked bars in **Figures S3A**, **S3B**). In black-headed gulls, inland colonies experienced a significant decline between 2022 and 2024 (estimated log difference = -0.44, SE = 0.14, $P = 0.011$), whereas no significant change was detected in coastal colonies overall. In common terns, inland colonies also showed a significant decline from 2022 to 2024 (estimated log difference = -0.43, SE = 0.13, $P = 8.3 \times 10^{-3}$), whereas coastal colonies remained relatively stable.

**Details on the Red List status of black-headed gull and common tern in Germany**

Assuming that our dataset captured nearly all breeding pairs of common terns in Germany in 2022 and subsequent years, the national population declined by more than 10% between 2016 and 2022, and by over 20% by 2024. In the 12 years prior to 2016, the common tern population had already declined by 6%, and by 9% since 1985, which classifies as a stable short-term and a negative long-term trend (Ryslavy et al., 2020). Aggregating these values suggests that common terns may have declined by 23–27% over the last 12 years, which is equivalent to an annual decrease of approximately 2%. According to the criteria of the German Red List of endangered avian species (Gerlach et al., 2019; Ryslavy et al., 2020), a breeding population decline of 1–3% per year over 12 years qualifies as a “Moderate Decline”. Given that the common tern now shows both a negative long-term and short-term trend, its current Red List status may warrant reassessment, potentially requiring a change from “Highly Threatened” to “Threatened with Extinction”. The black-headed gull remains a common breeding bird in Germany, with a long-term (-8% from 1980–2016) and short-term (+7% 2004–2016) stable breeding population (Gerlach et al., 2019; Ryslavy et al., 2020). It could remain classified as “Not Threatened”, despite the substantial reduction in breeding pair numbers of 15–24% within just two years.

### Supplementary figures

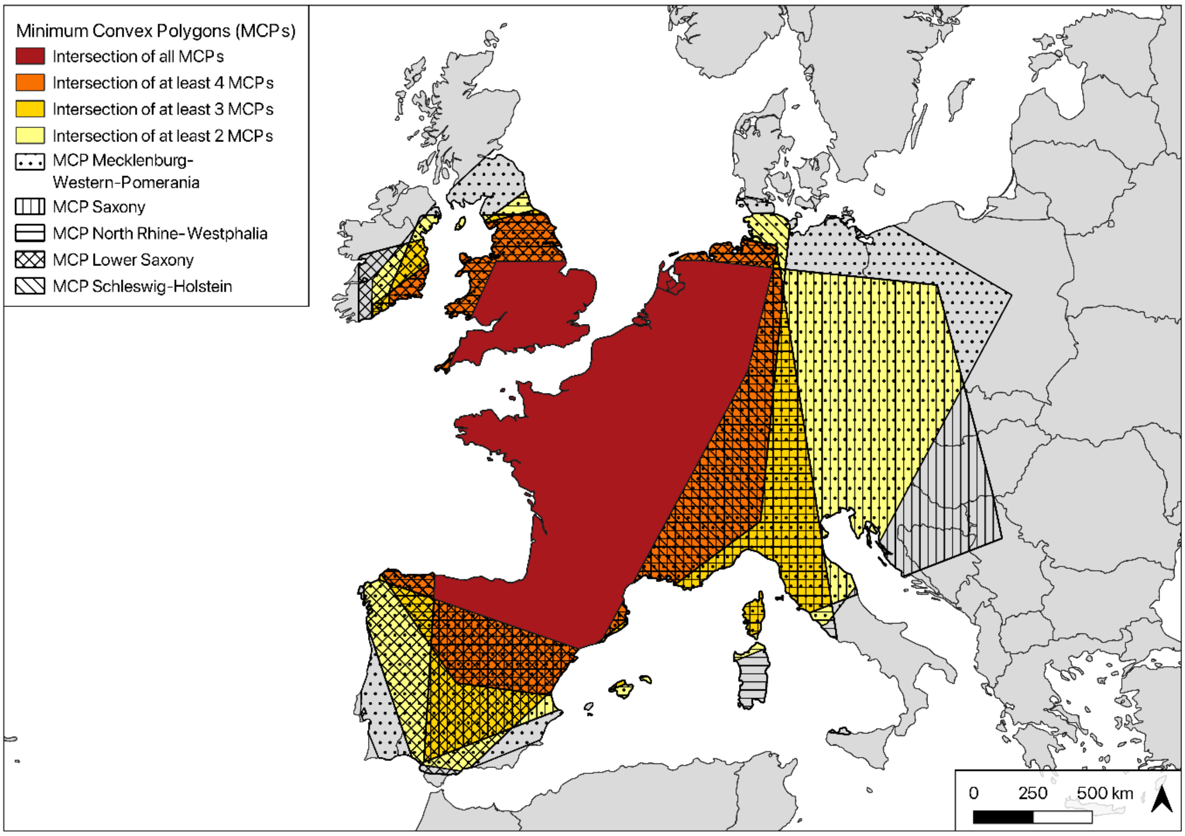


**Figure S1 |** Wintering areas of black-headed gulls according to the region (German federal state) where they had hatched or bred. MCP = minimum convex polygon, encompassing all winter recoveries of individuals originating from the respective regions.

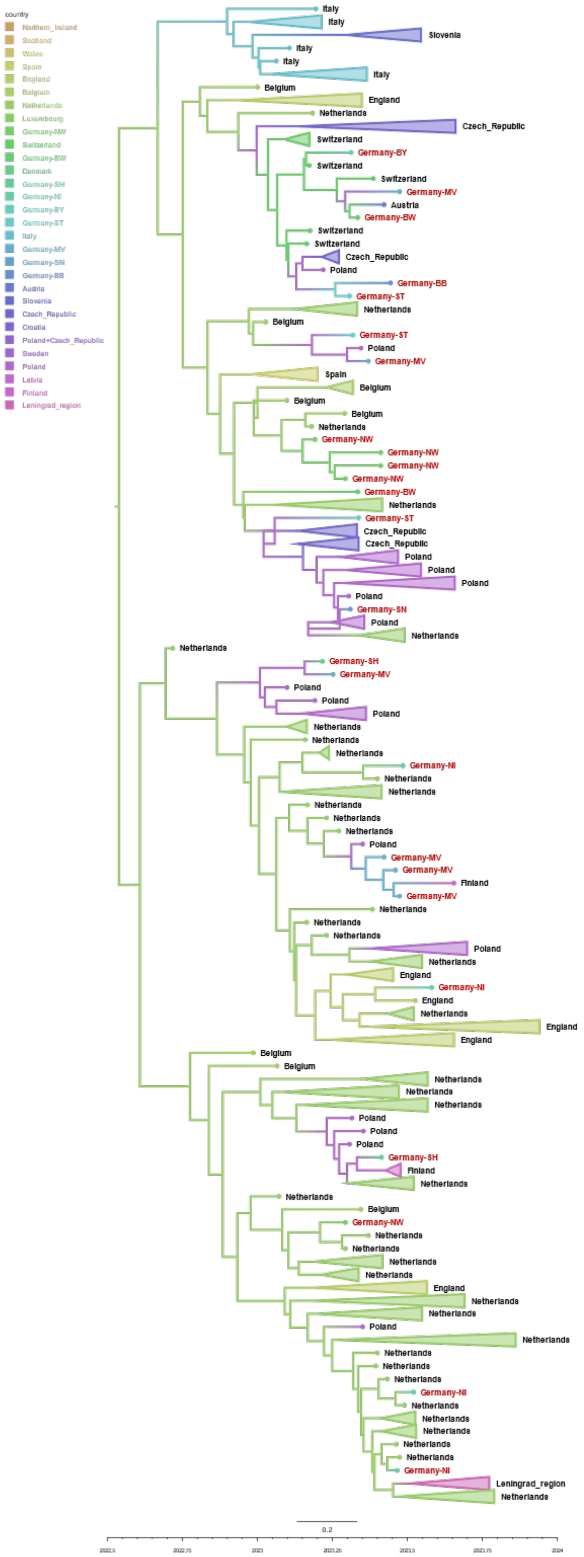


**Figure S2** | Time-scaled maximum clade credibility tree of highly pathogenic avian influenza virus H5N1 subtype 2.3.4.4b, genotype euBB, hemagglutinin coding sequences collected in Europe between September 2022 and December 2023. Branches are coloured by country of sampling, with countries ordered from west to east along a yellow-to-purple colour gradient. Tip labels of sequences originating from Germany are highlighted in red. Nodes not contributing to incursions into Germany have been collapsed for clarity.

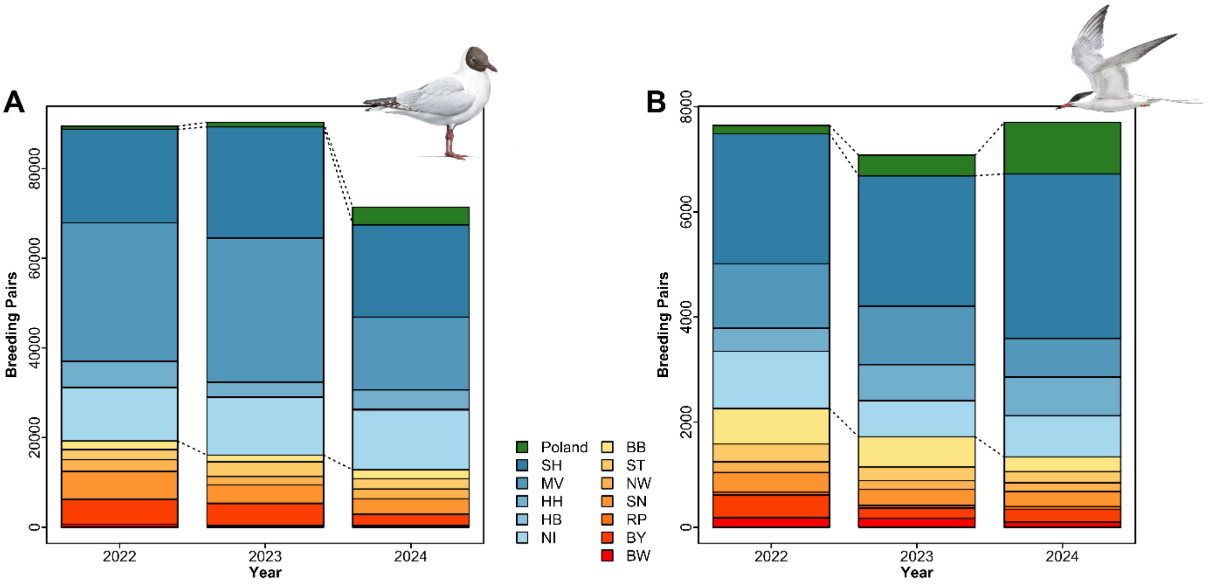


**Figure S3** | Changes in breeding pair numbers of (A) black-headed gulls and (B) common terns in Germany from 2022 to 2024. Breeding pair numbers from eastern Mecklenburg-Western Pomerania and the Szczecin Lagoon in Poland are included. Colours indicate inland (shades of red to yellow), coastal (shades of blue), and Polish (green) colonies. Note that the figure does not represent the entire German breeding population; data from inland Lower Saxony, and several other regions are missing. Relative to the most recent national census, our 2022 dataset represents 55.5–77.2% of the total number of breeding pairs in black-headed gulls and 83.2–88.1% in common terns. German federal states are shown with abbreviated names (SH: Schleswig-Holstein, MV: Mecklenburg-Western Pomerania, HH: Hamburg, HB: Bremen, NI: Lower Saxony, BB: Brandenburg, ST: Saxony-Anhalt, NW: North Rhine-Westphalia, SN: Saxony, RP: Rhineland-Palatinate, BY: Bavaria, BW: Baden-Württemberg).

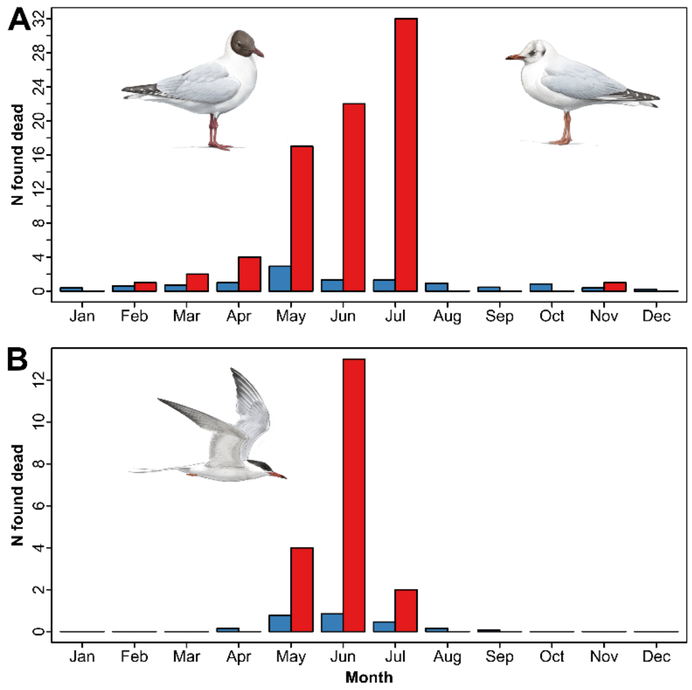


**Figure S4** | Reported monthly numbers of dead (A) black-headed gulls and (B) common terns found between 2010 and 2022 (monthly average, blue) and in 2023 (red) based on ring recoveries. All birds were ringed in eastern Germany. Note that the high number of dead black-headed gulls reported in July 2023 may be influenced by surveillance activity within the colonies: to minimize disturbances during the breeding season, site managers limited their presence in the colonies, meaning that birds found in July may have actually died in May or June.

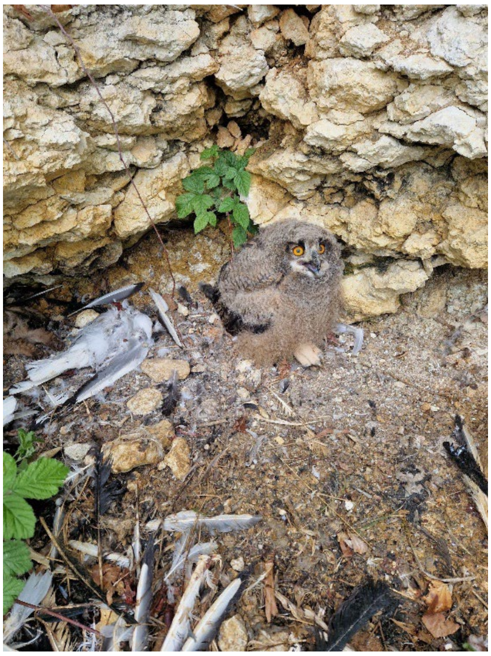

**Figure S5 |** Eagle owl (*Bubo bubo*) nestling with a black-headed gull carcass found in 2023 at the nest during ringing. Upon control a fortnight later, the nestling was found dead and tested positive for HPAIV H5N1 clade 2.3.4.4b, genotype euBB (Photo: Andreas Buck, used with permission from the Peregrine Falcon Protection Working Group Baden-Württemberg, 2023).

### Supplementary tables

**Table S1 |** Details on black-headed gulls and common terns tested for the presence of high pathogenicity avian influenza in the years 2022 to 2024, according to the German Avian-Influenza-Database (AI-DB). Infl A pos = Influenza A positive, H5 pos = H5 subtype-specific positive, HP = highly pathogenic positive.

| Species | Year | Quarter | *N* birds tested | Infl A pos (%) | H5 pos (%) | HP (%) |
|---|---|---|---|---|---|---|
| Black-headed gull | 2022 | I | 42 | 7 (16.7) | 6 (14.3) | 6 (14.3) |
| | | II | 32 | 14 (43.8) | 14 (43.8) | 14 (43.8) |
| | | III | 29 | 9 (31) | 9 (31) | 9 (31) |
| | | IV | 39 | 1 (2.6) | 1 (2.6) | 1 (2.6) |
| | 2023 | I | 118 | 73 (61.9) | 72 (61.0) | 52 (44.1) |
| | | II | 379 | 301 (79.5) | 254 (67.1) | 218 (57.5) |
| | | III | 54 | 39 (72.2) | 39 (72.2) | 33 (61.1) |
| | | IV | 26 | 4 (15.4)† | 2 (7.7) | 2 (7.7) |
| | 2024 | I | 25 | 2 (8.0)‡ | 1 (4.0) | 1 (4.0) |
| | | II | 15 | 0 (0.0) | 0 (0.0) | 0 (0.0) |
| | | III | 15 | 4 (26.7)‡ | 0 (0.0) | 0 (0.0) |
| | | IV | 30 | 0 (0.0) | 0 (0.0) | 0 (0.0) |
| Common tern | 2022 | I | 0 | — | — | — |
| | | II | 18 | 5 (20.8) | 5 (20.8) | 5 (20.8) |
| | | III | 26 | 6 (23.8) | 6 (23.8) | 6 (23.8) |
| | | IV | 0 | — | — | — |
| | 2023* | I | 0 | — | — | — |
| | | II | 133 | 99 (74.4) | 80 (60.2) | 43 (32.3) |
| | | III | 40 | 32 (80) | 21 (52.5) | 20 (50) |
| | | IV | 1 | 0 (0.0) | 0 (0.0) | 0 (0.0) |
| | 2024 | I | 0 | — | — | — |
| | | II | 3 | 0 (0.0) | 0 (0.0) | 0 (0.0) |
| | | III | 3 | 0 (0.0) | 0 (0.0) | 0 (0.0) |
| | | IV | 0 | — | — | — |

** In 2023, 6 HPAI H5-positive samples were taken from apparently healthy individuals.*

*† Two alive birds with different subtype H6N8.*

*‡ One bird in each time period with different subtype H16N3.*

**Table S2 |** Number of confirmed HPAIV cases in black-headed gulls, other gull species, and other wild birds during the winter months 2022/23 in Europe (data from TSN, ADIS and WOAH). Percentages refer to the total number of cases reported in the entire time period.

| Species | November | December | January | February |
|---|---|---|---|---|
| Black-headed gull | 5 (0.27%) | 5 (0.27%) | 77 (4.20%) | 336 (18.31%) |
| Unspecified or other gull species | 23 (1.25%) | 14 (0.76%) | 44 (2.40%) | 98 (5.34%) |
| Other wild bird species | 354 (19.29%) | 226 (12.31%) | 329 (17.93%) | 324 (17.66%) |
| Total | 382 (20.82%) | 245 (13.35%) | 450 (24.52%) | 758 (41.31%) |

**Table S3** | Descriptive statistics for the presence of influenza A specific antibodies in adult breeding black-headed gulls and common terns, as well as common tern chicks sampled on day 20 post-hatching at the Isle of Böhmke or Banter See during the 2022–2024 breeding seasons. The table shows the total numbers of individuals sampled and those testing seropositive for general influenza A virus (IAV) antibodies or for H5 subtype-specific antibodies.

| **Year** | **Place** | **Species** | **Age** | ***N*** | **IAV (%)** | **H5 (%)** |
|---|---|---|---|---|---|---|
| 2022 | Banter See | Common tern | adult | 152 | 27 (17.8) | * |
| 2023 | Isle of Böhmke | Black-headed gull | adult | 22 | 15 (68.2) | 1 (6.7) |
| | Banter See | Common tern | adult | 253 | 30 (11.9) | 3 (10.0) |
| | Banter See | Common tern | chick d20 | 12 | 0 (0.00) | — |
| 2024 | Banter See | Common tern | adult | 168 | 68 (40.5) | 23 (34.0) |
| | Banter See | Common tern | chick d20 | 143 | 2 (1.40) | 0 (0.00) |

** In the year of the first H5N1 outbreak (2022), insufficient serum volumes prevented H5 subtype-specific testing in the 27 adult common terns that tested positive for general IAV antibodies.*

### Supplementary acknowledgments

We are deeply grateful to all those who provided mortality data from their respective black-headed gull and common tern colonies: Phillip Aldinger (Landkreis Nordwestmecklenburg); BUND Naturschutzzentrum Möggingen; Elmar Ballstaedt (Verein Jordsand zum Schutz der Seevögel und der Natur e.V.); Martin Beyer (Landkreis Vorpommern-Greifswald); Claudius Birke (Landesbund für Vogel- und Naturschutz in Bayern e.V. [LBV]); Christian Brummer (LBV); Holger Bruns (Naturschutzbund Deutschland e.V. [NABU]); Norman Donner (Nationalpark Vorpommersche Boddenlandschaft); Helene Falk (Schutzgemeinschaft Ammersee e.V.); Bastian Forkel (LBV); Silke Fregin (Fachgruppe Ornithologie Greifswald); Katrin Fritzsch (NABU); Karl Fidelis Gauggel (NABU); Wolfgang Gaus (GAU Schutzgemeinschaft für den Neu-Ulmer Lebensraum e.V.); Andrea Gehrold (LBV); Felix Gruetzmacher; Thomas Heinicke (Arbeitsgemeinschaft Berlin-Brandenburgischer Ornithologen); Bernd Heinze (Landesamt für Umwelt, Naturschutz und Geologie Mecklenburg-Vorpommern); Veit Hennig (Verein Jordsand zum Schutz der Seevögel und der Natur e.V.); Christian Huber (LBV); Frank Joisten; Stella Klasan (Landkreis Vorpommern-Greifswald); Thomas Klinner (Naturwacht Brandenburg); Steffen Koschkar; Clemens Krafft (Schutzgemeinschaft Ammersee e.V.); Brigitte Kraft (LBV); Patrick Kretz (Nordrhein-Westfälische Ornithologengesellschaft e.V.); Thomas Krämer (Biologischen Station Rieselfelder Münster); Bernd Litzkow; Sebastian Lorenz (Universität Greifswald); Lisa Maier (NABU); Helma Mensing (Biologische Station Zwillbrock e.V.); Norbert Model (LBV); Mathias Mähler (Thüringer Landesamt für Umwelt, Bergbau und Naturschutz); Reinhardt Möckel; Abbo van Neer (Kreis Dithmarschen); Walter Niederer (Naturschutzverein Rheindelta); Florian Packmor (Nationalpark Niedersächsisches Wattenmeer); Alfons Pennekamp; Raphael Rehm (ARGE Schwäbisches Donaumoos e.V.); Richard Roberts; Martin von Roeder (Gesellschaft für Naturschutz und Ornithologie Rheinland-Pfalz e.V.); Carolin Rothfuß (Verein Jordsand zum Schutz der Seevögel und der Natur e.V.); Martin Sansoni (Landkreis Straubing-Bogen); Helmut Schmitt; Fabian Sieg (Biosphärenreservat Mittelelbe); Wilfried Starke (Fachgruppe Ornithologie Greifswald); Ole Stejskal (Niedersächsisches Landesamt für Verbraucherschutz und Lebensmittelsicherheit); Helmut Stocker; Stefan Sudmann (Planungsbüro STERNA); Aleksandra Szwagierczak (LBV); Jan Tenner (Unteren Naturschutzbehörde in Neuburg-Schrobenhausen); Peter Thiele; René Thiemann (Landesamt für Umweltschutz Sachsen-Anhalt); Hendrik Trapp (Sächsisches Landesamt für Umwelt, Landwirtschaft und Geologie); Veterinäramt Landkreis Leipzig; Veterinäramt Stadt Leipzig; Frank Vökler (Ornithologische Arbeitsgemeinschaft Mecklenburg-Vorpommern e.V.); Georg Walcher; Josef Yun (Staatliches Veterinäramt Landshut); Peter Zach (LBV); and the many additional volunteers whose contributions were compiled by those named above.

Equally, we extend our thanks to the following individuals and institutions for providing data on breeding pair numbers of both species: the Federation of German Avifaunists (Dachverband Deutscher Avifaunisten e.V., DDA); Birger Reibisch (Ornithologische Arbeitsgemeinschaft für Schleswig-Holstein und Hamburg e.V., SH); Leonie Enners and Bernd Hälterlein (Nationalparkverwaltung Schleswig-Holsteinisches Wattenmeer, SH); Markus Risch (Bündnis Naturschutz in Dithmarschen e.V., SH); Sven Baumung (Behörde für Umwelt, Klima, Energie und Agrarwirtschaft, HH); Florian Packmor (Nationalparkverwaltung Niedersächsisches Wattenmeer, NI); Torsten Ryslavy (Landesamt für Umwelt Brandenburg, BB); Stefan Fischer (Landesamt für Umweltschutz Sachsen-Anhalt, ST); Stefan Sudmann (Planungsbüro STERNA, NW); Alfons Pennekamp (NW); Hendrik Trapp (Sächsisches Landesamt für Umwelt, Landwirtschaft und Geologie, SN); Martin von Roeder (Gesellschaft für Naturschutz und Ornithologie Rheinland-Pfalz e.V., RP); Christian Brummer, Alexandra Fink, Bastian Forkel, Andrea Gehrold, and Peter Zach (all LBV, BY); Wolfgang Gaus (GAU Schutzgemeinschaft für den Neu-Ulmer Lebensraum e.V., BY); Max Kurzmann (OAG Chiemsee, BY); Jost Einstein (Ornithologische Gesellschaft Baden-Württemberg, BW); Katrin Fritzsch (NABU Federsee, BW); Lisa Maier (NABU Bodensee, BW); and the many additional volunteers whose contributions were compiled by those named above.

We are also grateful to all bird ringers who contributed data via the three German ringing schemes: the Vogelwarten Hiddensee, Helgoland, and Radolfzell.

We also thank the city of Wilhelmshaven for continued access to the common tern colony at Banter See, as well as Nathalie Kürten, Maria Moiron, Benno Rodemann, Johanna Eilers and various students and volunteers for the heartbreaking work of daily collection of dead terns during the seasons of 2022 and 2023.

We are grateful to Javier Lazaro for his black-headed gull and common tern illustrations and to Andreas Buck and the Peregrine Falcon Protection Working Group Baden-Württemberg for providing the photo of the Eagle owl nestling.

### ORCID references of all authors

Ulrich Knief: 0000-0001-6959-3033
Ann Kathrin Ahrens: 0009-0007-3635-423X
Valerie Allendorf: 0000-0002-8703-1840
Justine Bertram: 0000-0003-3364-9570
Sandra Bouwhuis: 0000-0003-4023-1578
Wolfgang Fiedler: 0000-0003-1082-4161
Anja Globig: 0009-0006-1966-6760
Anne Günther: 0000-0001-5176-6346
Christof Herrmann: 0009-0008-4497-3345
Sascha Knauf: 0000-0001-5744-4946
Dominik Marchowski: 0000-0001-7508-9466
Simon Piro: 0000-0002-1035-4246
Anne Pohlmann: 0000-0002-5318-665X
Robert E. Rollins: 0000-0002-5779-7001
Christoph Staubach: 0000-0002-1574-0138
Timm Harder: 0000-0003-2387-378X
